# *Cannabis sativa* Terpenes are Cannabimimetic and Provide Support for the Entourage Effect Hypothesis

**DOI:** 10.1101/2020.10.22.350868

**Authors:** Justin E. LaVigne, Ryan Hecksel, Attila Keresztes, John M. Streicher

**Affiliations:** Department of Pharmacology, College of Medicine, University of Arizona, Tucson, AZ USA

**Author notes:** Author Contributions J.E.L. designed research, performed research, analyzed data, wrote paper. R.H. performed research, analyzed data, edited paper. A.K. performed research, analyzed data, edited paper. J.M.S. analyzed data, edited paper.

**Keywords:** Cannabis, terpene, entourage effect, cannabinoid

## Abstract

Limited evidence has suggested that terpenes found in *Cannabis sativa* are analgesic, and could produce an “entourage effect” whereby they modulate cannabinoids to result in improved outcomes. However this hypothesis is controversial, with limited evidence. We thus investigated *Cannabis sativa* terpenes alone and with the cannabinoid agonist WIN55,212 using *in vitro* and *in vivo* approaches. We found that the terpenes α-humulene, geraniol, linalool, and β-pinene produced cannabinoid tetrad behaviors in mice, suggesting cannabimimetic activity. Some behaviors could be blocked by cannabinoid or adenosine receptor antagonists, suggesting a mixed mechanism of action. These behavioral effects were additive with WIN55,212, providing support for a terpene “entourage effect.” *In vitro* experiments showed that all terpenes activated the CB1R, while some activated other targets. Our findings suggest that these *Cannabis* terpenes are multifunctional cannabimimetic ligands that provide support for the entourage effect hypothesis and could be used to enhance the therapeutic properties of cannabinoids.

## Introduction

*Cannabis sativa* is a dioecious plant belonging to the Cannabaceae family, along with another popular plant, *Humulus lupulus* (hops) [1]. The plant itself is a “biopharmacy” containing hundreds of phytochemicals [2], many with medicinal indications. Of these, the phytocannabinoids and terpenes have been the most studied in regard to their medicinal and therapeutic properties [3, 4]. Terpenes, which are the basic constituents of essential oils found in many plants, have been used for thousands of years for therapeutic purposes. They also provide flavor and aroma for cannabis and other plants. Studies in animal models and humans have identified analgesic, anti-microbial, anti-inflammatory, and similar therapeutic properties for terpene treatment [5-7]. Phyto-cannabinoids, most notably Δ9-tetrahydrocannabinol (THC), have been the main focus of research for mechanistic and therapeutic studies [4]. While cannabis contains both of these families of phytochemicals, the terpenes have been less studied than the phytocannabinoids, and the potential interaction between terpenes and phytocannabinoids when the plant is consumed for recreational and medicinal purposes has barely been studied at all.

The hypothesized interactions between various phytocannabinoids and terpenes to produce unique outcomes from either chemical alone is known as the “entourage effect” [3, 5, 8]. The evidence for the entourage effect is comprised of deductive reasoning arguments [5, 9], some clinical suggestions [10, 11], and a few pre-clinical investigations [8, 12-14]. There is also skepticism within the literature [15], and some evidence against cannabinoid and terpene interactions from preclinical studies [7, 16]. It remains unclear whether terpenes can influence the activity of cannabinoids, and if they do, whether this modulation is a result of direct influence on cannabinoid receptors, as with β-caryophyllene [13], or indirect modulation via other mechanisms.

If the entourage effect can be demonstrated, it could provide a powerful tool to improve cannabinoid therapy. The main phytocannabinoids THC and cannabidiol (CBD) work through cannabinoid and non-cannabinoid mechanisms [4, 17] to evoke therapeutic benefits, most notably treatment for chronic pain [18, 19]. However efficacy in these studies tends to be modest, and THC induces burdensome psychoactive and somatic side effects [19-21]. If terpene compounds modulate phytocannabinoids like THC, then it might be possible to identify terpenes that maximize the therapeutic efficacy of cannabinoids while reducing unwanted side effects. Therapeutically, this could take the form of specific chemovar plant strains, or purified and defined terpene/cannabinoid mixtures.

In this study we thus aimed to assess whether the entourage effect is a *possible* phenomenon by observing the actions of various terpenes *in vivo* and *in vitro* both alone and in combination with an established cannabinoid agonist. Our results establish a *possible* entourage effect between cannabinoids and terpenes by demonstrating that selected terpenes have poly-pharmacological effects at both cannabinoid and non-cannabinoid receptors, and selectively modulate canonical cannabinoid agonist activity.

## Results

### Terpenes Induce Cannabinoid Tetrad Behaviors in Mice

Our first set of experiments sought to determine whether selected terpenes (α-Humulene, β-Pinene, Linalool, Geraniol, and β-Caryophyllene as a putative cannabinoid receptor type 2 [CB2] agonist control) had activity in the cannabinoid tetrad of behaviors mediated by the CB1 receptor: antinociception, hypolocomotion, catalepsy, and hypothermia [20]. These terpenes were selected based on their quantities in *Cannabis sativa*, with α-Humulene, β-Pinene, Linalool, and β-Caryophyllene all being found in higher quantities and Geraniol in lower to underdetermined quantities. Terpenes were administered at several doses (50mg/kg – 200mg/kg) *i*.*p*. and assessed in the tail flick assay in male and female CD-1 mice. As β-Caryophyllene has been identified as a selective CB2 agonist, and induced CB2-mediated effects at 50mg/kg [13], we administered it at 100mg/kg as a known CB2 agonist for our behavioral assays. Thus, if selective, it should not exhibit the distinct CB1-mediated tetrad behaviors.

Terpenes induced a range of efficacies in the tail flick assay **(Figure S1A-E)**. Geraniol and α-Humulene exhibited moderate ∼40-50% efficacy in a dose-dependent manner **(Figure S1A and S1D)** while β-Pinene showed low efficacy but not in a dose-dependent manner **(Figure S1B)**, suggesting partial agonist activity at the top of the dose range. Linalool demonstrated dose-dependent low efficacy **(Figure S1C)**, as did the β-Caryophyllene at the dose tested **(Figure S1E)**. The positive control WIN55,212-2 demonstrated dose-dependent increases in thermal latency with greater efficacy than any of the tested compounds reaching near-threshold values at a 10 mg/kg dose **(Figure S1F)**.

Terpenes were next assessed in the remaining tetrad behaviors during the peak effect window observed in the tail flick assay (i.e. 0-30 min post-injection). α-Humulene, β-Pinene, Geraniol and Linalool, as well as the control WIN55,212-2, induced significant hypothermia **(Figure S2A)**, significant increases in cataleptic behavior **(Figure S2B)**, and significant reductions in locomotor activity **(Figure S2C and S2D)** compared to their baseline values. The CB2-selective control, β-Caryophyllene, as well as Vehicle control did not induce hypothermia and catalepsy. Hypolocomotion observations were likely confounded by mouse habituation of the open field boxes between baseline and post-injection measurements, as shown in Vehicle treatment in **Figure S2C and S2D**. Therefore, the experiment was repeated without a baseline recording. In this experiment WIN55,212-2, β-Pinene, Geraniol and Linalool all displayed reductions in distance traveled and mobile time, whereas Vehicle, β-Caryophyllene or α-Humulene treatment did not result in significant reductions **(Figure S3A, S3B)**. However, α-Humulene did trend towards significance (p = 0.057). Together these results suggest that our terpenes are cannabimimetic, inducing at least 3 out of 4 of the classic cannabinoid tetrad behaviors each, while Vehicle and β-Caryophyllene controls did not. A radar chart depiction of the impact of the different terpenes on tetrad behavior is shown in **Figure 1**.

**Figure 1:**
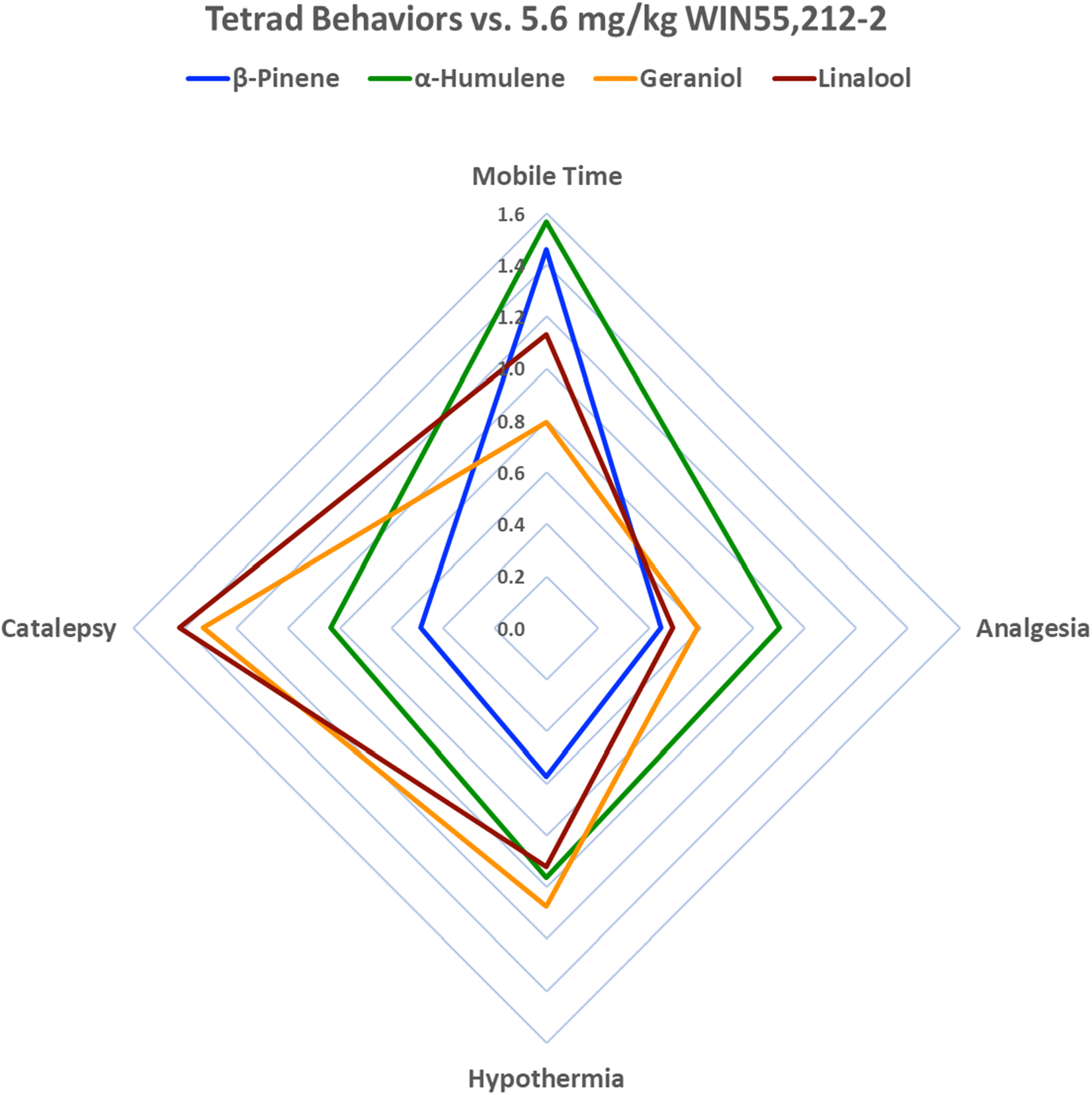
Terpenes Induce Cannabimimetic Tetrad Behaviors in Mice. Behavioral analysis data from **Figures S1-S3** represented in a radar chart format. The percent change from baseline for each terpene in each behavior was determined; the peak effect for tail flick was selected (**Figure S1**) while the other behaviors only had a single time point. This percent change was then normalized to the % change of 5.6 mg/kg WIN55,212-2 (1.0 on the chart above). The chart demonstrates that while all terpenes induce all 4 tetrad behaviors, they do so with varying efficacy; they also vary whether the effect is stronger or weaker than the positive control WIN55,212-2.

### Terpene Tail Flick Antinociception is CB1 Mediated and is Additive with Cannabinoid

To determine the role of the CB1 receptor in potentially mediating these tetrad behaviors induced by terpenes we used the CB1 selective antagonist/inverse agonist rimonabant. We first showed that rimonabant could fully or partially reverse the tetrad behaviors induced by the positive control cannabinoid WIN55,212-2 (**Figure S4**). We then used this drug in terpene tail flick anti-nociception, and showed that rimonabant pretreatment fully blocked terpene response in this assay, suggesting that terpenes induce tail flick anti-nociception via the CB1 **(Figure 2)**.

**Figure 2:**
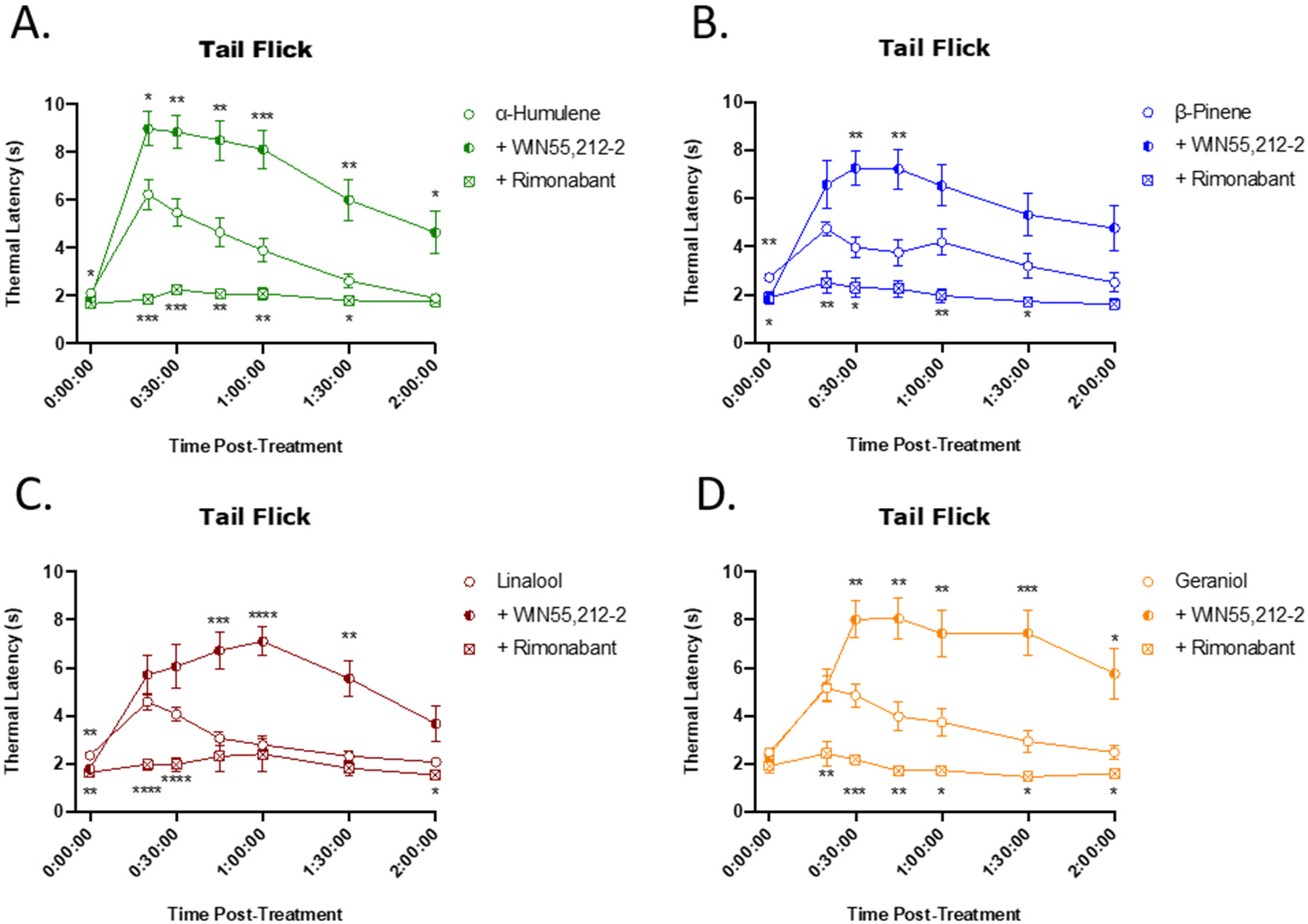
Terpenes Induce CB1 Dependent and Cannabinoid Additive Antinociception in the Tail Flick Assay. Mice were treated with 200mg/kg terpene alone, combined with 5.6 mg/kg WIN55,212-2, or after pretreatment with 10 mg/kg rimonabant, all injected *i*.*p*.. Mice were then assessed in the tail flick test over 2 hr. **A)** α-Humulene, **B)** β-Pinene, **C)** Linalool and **D)** Geraniol. Data represents the mean ± SEM of tail flick latency (n=10-14/group). Statistics analyzed via RM two-way ANOVA, Dunnett’s *post hoc*; *p<0.05, ** p<0.01, *** p<0.001, **** p<0.0001, compared to terpene alone at same time point.

In order to test potential terpene/cannabinoid interaction, terpene was combined with WIN55,212-2 in the tail flick assay. When a given terpene was combined with a lower dose of WIN55,212-2, the combined effect was increased compared to terpene or WIN55,212-2 alone **(Figure 2)**, demonstrating a terpene/cannabinoid interaction in modulating antinociception. Whether this interaction is additive or synergistic in nature is currently under investigation. This observation lends potential support to the “entourage effect” hypothesis.

However, as an inverse agonist, rimonabant has the potential to reverse tail flick responses through a systems level inverse agonism effect on antinociception. In other words, rimonabant could demonstrate blockade of antinociception through pro-nociceptive activity. To control for this, we assessed the effect of rimonabant pretreatment on morphine-induced antinociception in the tail flick assay **(Figure S5A-B)**, at an equal-efficacy dose compared to our terpenes, and on baseline thermal latency responses at a reduced water bath temperature **(Figure S5C)**. Rimonabant had no significant effect on morphine-induced antinociception or baseline thermal latency responses, further suggesting terpenes mediate their antinociceptive actions through a CB1-dependent mechanism.

### Terpene Hypothermia is Additive with Cannabinoid but Mostly Not Mediated by CB1

Following the experimental design used for tail flick antinociception above, we next sought to determine the mechanisms of terpene-induced hypothermia. First, when we co-injected both terpene and WIN55,212-2, hypothermia was increased over either treatment alone for all terpenes tested (**Figure 3**). This is similar to the finding for tail flick anti-nociception, and further lends support to the entourage effect hypothesis.

**Figure 3:**
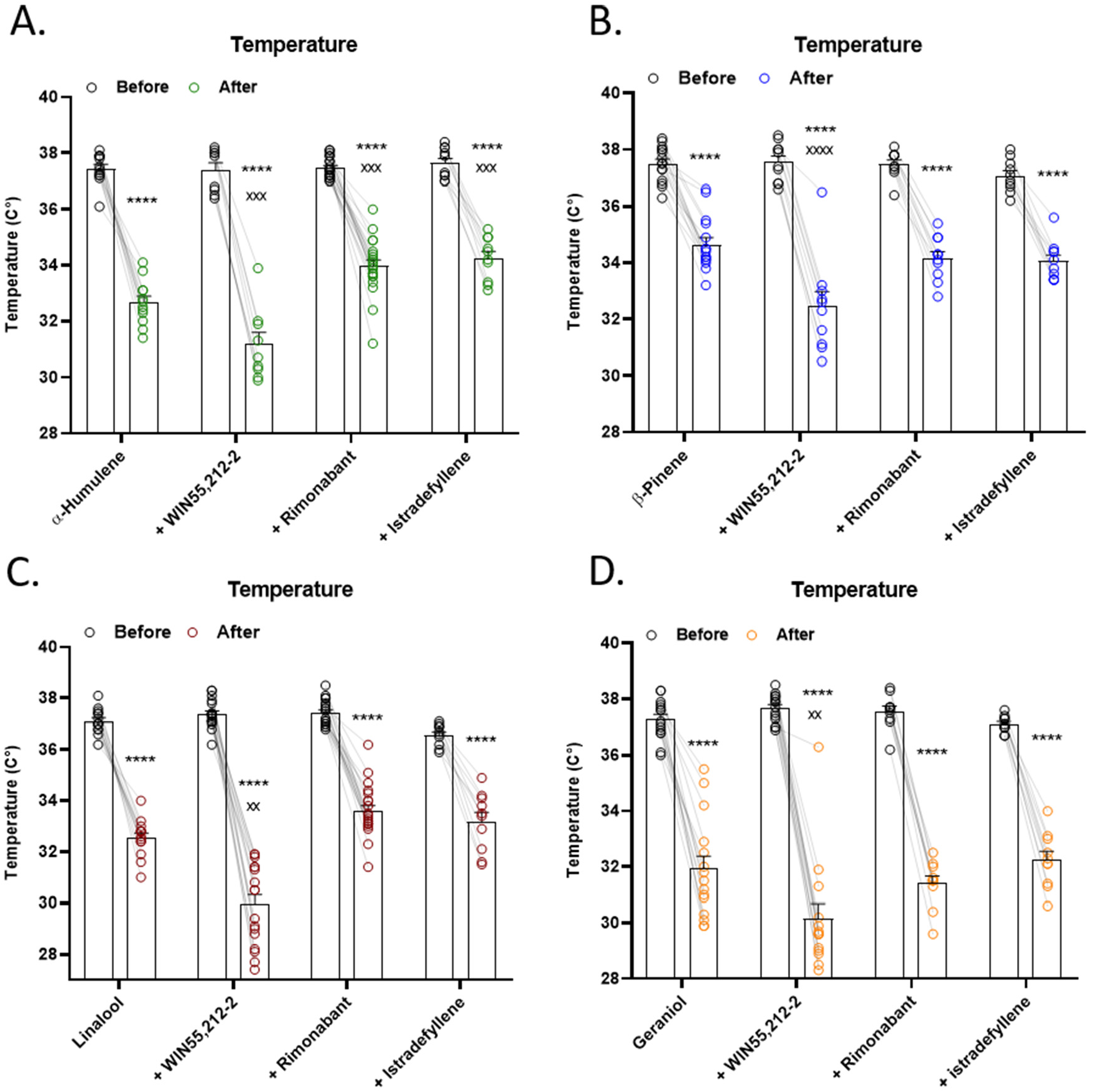
Terpenes Induce Hypothermia Through Mostly Non-CB1/A2a Mechanisms and are Additive with Cannabinoid. Mice were baselined for temperature, injected with 200 mg/kg terpene alone, combined with 5.6 mg/kg WIN55,212-2, or after pretreatment with 10 mg/kg rimonabant or 10 mg/kg istradefyllene. After 30min, temperature was assessed again. **A)** α-Humulene, **B)** β-Pinene, **C)** Linalool and **D)** Geraniol. Data represents the mean ± SEM temperature (n=10-20/group). Statistics analyzed via RM two-way ANOVA, Tukey’s *post hoc*; **** p<0.0001 compared to each baseline; xx p<0.01, xxx p<0.001, xxxx p<0.0001 compared to terpene post-treatment.

However, unlike for tail flick, rimonabant treatment was only able to partially reverse α-humulene hypothermia (**Figure 3A**), and had no effect on the other terpenes. While as shown above rimonabant only partially reversed WIN55,212-2 hypothermia (**Figure S4C**), this still suggests that CB1 may only mediate α-humulene hypothermia, and has no role for the other terpenes. This further suggests that while the terpenes are cannabimimetic, they may induce these effects through both CB1-dependent and independent mechanisms.

Seeking to test the involvement of other receptor systems, we tested the role of adenosine A2a receptors (A2a) in terpene-induced hypothermia. Activation of A2a can induce hypothermia, hypolocomotion and cataleptic-like behaviors [22] and many studies have started to investigate the interactions between the cannabinoid and adenosine systems [23-27]. In fact, a known cannabis terpene, D-limonene, is an agonist at the A2a receptor [28]. We thus hypothesized that activation of A2a receptors may contribute to the rimonabant-insensitive behaviors induced by the studied terpenes. However, pretreatment with the A2a antagonist istradefyllene partially reversed α-humulene hypothermia (**Figure 3A**) while having no effect on the other terpenes, suggesting no role for the A2a much like the CB1 for terpene-induced hypothermia. Further complicating the analysis of the A2a in our behaviors, we found that istradefyllene alone had a small but significant hypothermic and hyperlocomotive effect, suggesting these behaviors be carefully interpreted (**Figure S6**).

### Terpene Catalepsy is Partially Additive with Cannabinoid and Mostly Mediated by A2a

We continued our mechanistic analysis with terpene-induced catalepsy. In contrast to the above findings, combining terpene with WIN55,212-2 did produce additive effects with α-humulene and β-pinene (**Figure 4A-B**), but not Linalool and Geraniol (**Figure 4C-D**). Here we begin to see differentiation of our “entourage effect” evidence, which suggests that different terpenes could be used to modulate different parts of the cannabinoid response. We also found that rimonabant had no impact on the catalepsy response for any terpene, suggesting this terpene behavior is CB1-independent (**Figure 4**).

**Figure 4:**
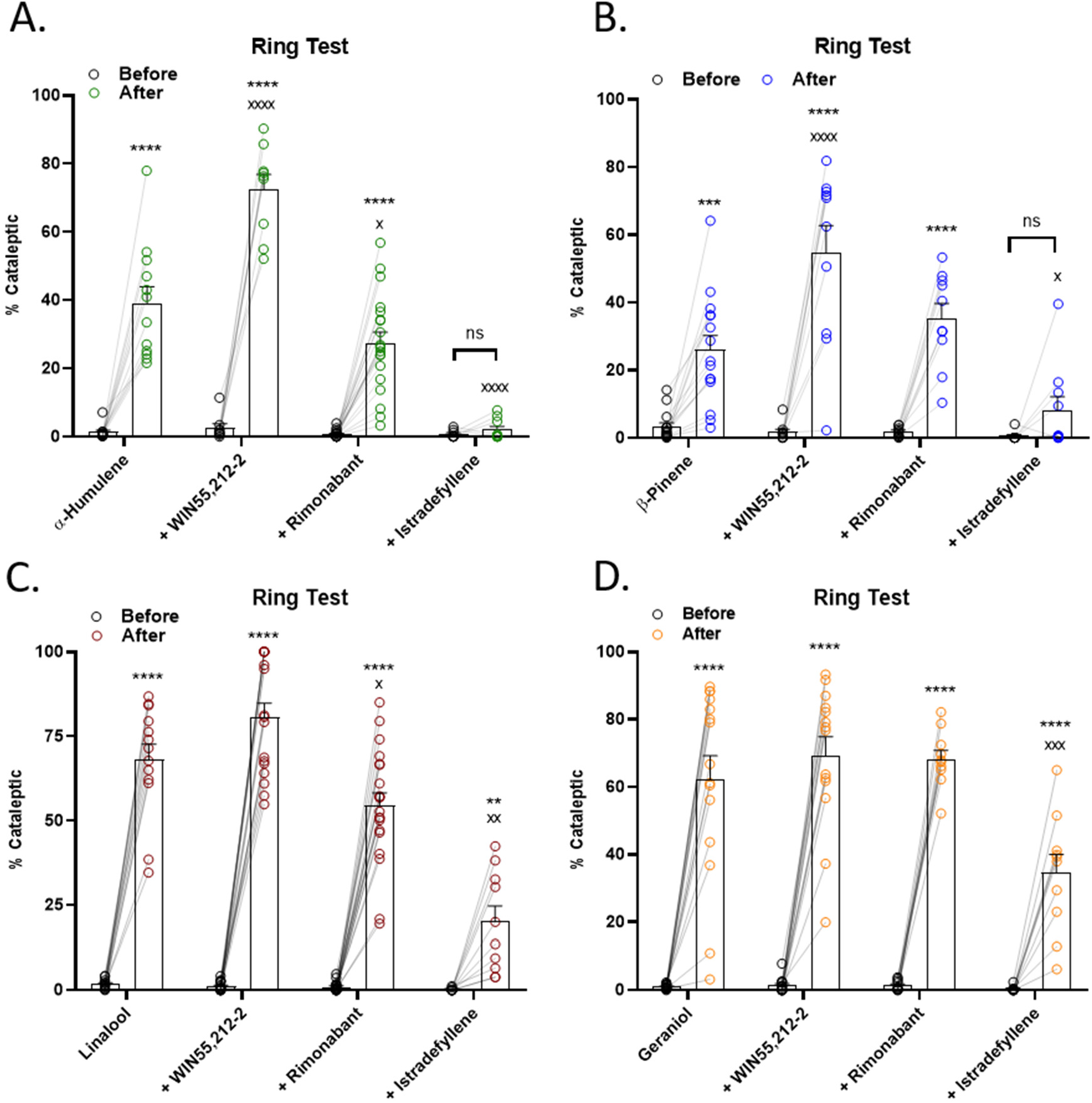
Terpene Induced Catalepsy is Partially Mediated by CB1, Strongly Mediated by A2a, and is Partially Additive with Cannabinoid. Mice were baselined in the ring test for 5 min, injected with 200 mg/kg terpene alone, combined with 5.6 mg/kg WIN55,212-2, or after pretreatment with 10 mg/kg rimonabant or 10 mg/kg istradefyllene. After 15 min, mice were tested in the ring test again for 5 min. **A)** α-Humulene, **B)** β-Pinene, **C)** Linalool and **D)** Geraniol. Data represents the mean ± SEM of % catalepsy (n=10-20/group). Statistics analyzed via RM two-way ANOVA, Tukey’s *post hoc*; ** p<0.01, *** p<0.001, **** p<0.0001, not significant (ns) compared to baseline; x p<0.05, xx p<0.01, xxx p<0.001, xxxx p<0.0001 compared to terpene post-treatment.

However, we did find a major role for the A2a in this behavior. Istradefyllene pretreatment completely blocked the catalepsy response for α-humulene and β-pinene, suggesting that the A2a is a necessary component of catalepsy for these terpenes **(Figure 4A-B**). Istradefyllene also partially blocked catalepsy for Linalool and Geraniol, showing that it is still a significant part of the mechanism for these terpenes (**Figure 4C-D**). Our results thus suggest that the A2a has a major role in terpene-induced catalepsy, but not the CB1.

### Terpene Hypolocomotion is Partially Additive with Cannabinoid and Partially A2a Mediated

Our analysis of mobile time as a measure of hypolocomotion is shown in **Figure 5**. We did observe a further decrease in mobile time when terpene was combined with WIN55,212-2 for α-humulene, β-pinene, and Linalool (**Figure 5A-C**) but not Geraniol (**Figure 5D**). This continues our theme of generally showing terpene/cannabinoid additive effects, albeit with specific behavioral impacts for each terpene. Also of note, rimonabant had no reversal effect on mobile time for any terpene, suggesting this behavior is also CB1-independent. However, we did observe significant reversal with istradefyllene for β-pinene (**Figure 5B**) and Geraniol (**Figure 5D**), showing that the A2a is mediating this behavior for some terpenes. We also report the distance traveled measurement of hypolocomotion in **Figure S7**; this data was in general less robust than the mobile time, although we did observe istradefyllene reversal with β-pinene (**Figure S7B**) and Geraniol (**Figure S7D**), confirming this finding from **Figure 5**.

**Figure 5:**
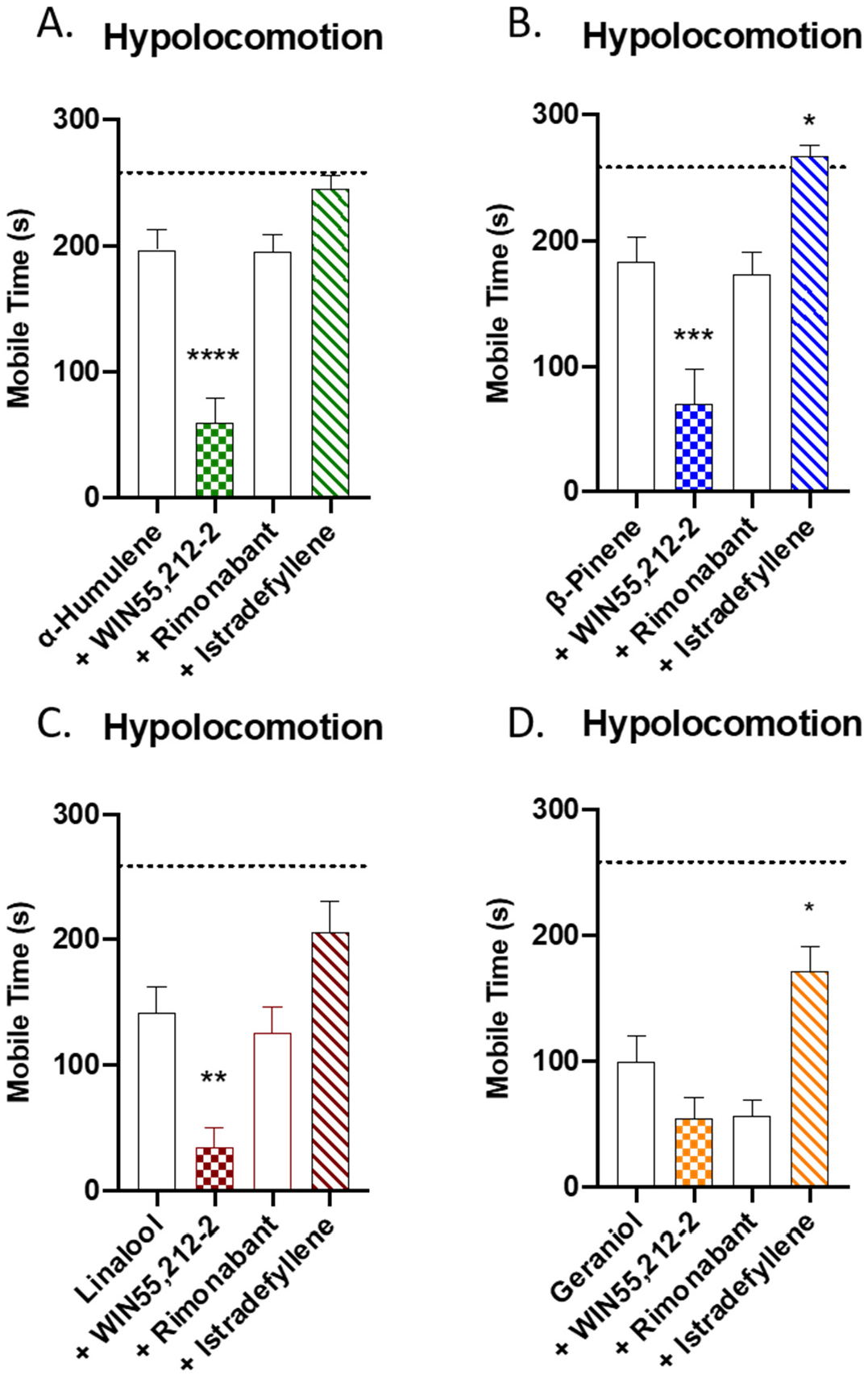
Terpene Induced Hypolocomotion is Partially Mediated by A2a and is Additive with Cannabinoid. Mice were injected with 200 mg/kg terpene alone, combined with 5.6mg/kg WIN55,212-2, or after pretreatment with 10 mg/kg rimonabant or 10 mg/kg istradefyllene, *i*.*p*.. After 10 min mice were then placed back into the open field box for a 5 min test. **A)** α-Humulene, **B)** β-Pinene, **C)** Linalool and **D)** Geraniol. Data represents the mean ± SEM of mobile time seconds (n=10-20/group). Statistics analyzed via one-way ANOVA, Dunnett’s *post hoc*; * p<0.05, ** p<0.01, *** p<0.001, **** p <0.0001 compared to terpene alone. Dotted line denotes vehicle levels of mobile time for reference.

Overall, our mechanistic studies suggest that the terpenes tested generally although selectively increase the behavioral effects of the cannabinoid WIN55,212-2, supporting the entourage effect hypothesis. Our findings also suggest a mix of CB1-dependent, A2a-dependent, and independent mechanisms for terpene behavioral effects. These findings overall suggest that *Cannabis* terpenes can have significant pharmacological effects, and could be used to selectively modulate the impact of *Cannabis*/cannabinoid therapy.

### Linalool has Specific Sex Differences

For all experiments above, both male and female mice were used, and in nearly every case, no differences were observed. However, we did observe specific sex differences for Linalool. First, in the tail flick assay, both males and females had the same response to Linalool alone, and both were fully blocked by rimonabant. However, we observed a sex difference when Linalool was combined with WIN55,212-2; males showed greater additive effects of combining both that occurred earlier in the time course, while females showed a delay in response and no potentiation over Linalool alone (**Figure S8A**). For Linalool hypolocomotion, we also observed a mechanistic difference; males showed rimonabant reversal of the behavior while females showed istradefyllene reversal, suggesting this behavior is mediated by CB1 in males and A2a in females (**Figure S8B-C**). These observations suggest that other terpene sex differences could be found, although the mechanism for this difference is unknown.

### Terpenes Activate the CB1 In Vitro

Our behavioral results suggested that the terpenes potentially interact with the CB1 receptor, and likely others. We thus sought to determine whether these selected terpenes acted as CB1 receptor agonists *in vitro*, first by assessing CB1-dependent ERK activation. Each of the terpenes tested, including the putative CB2 agonist β-Caryophyllene, activated downstream ERK signaling in CB1-CHO cells **(Figures 6, S9)**. This activation was rimonabant-sensitive **(Figures 7A, S10A-B)**, suggesting that terpenes act as CB1 agonists *in vitro*, further supporting our *in vivo* findings.

**Figure 6:**
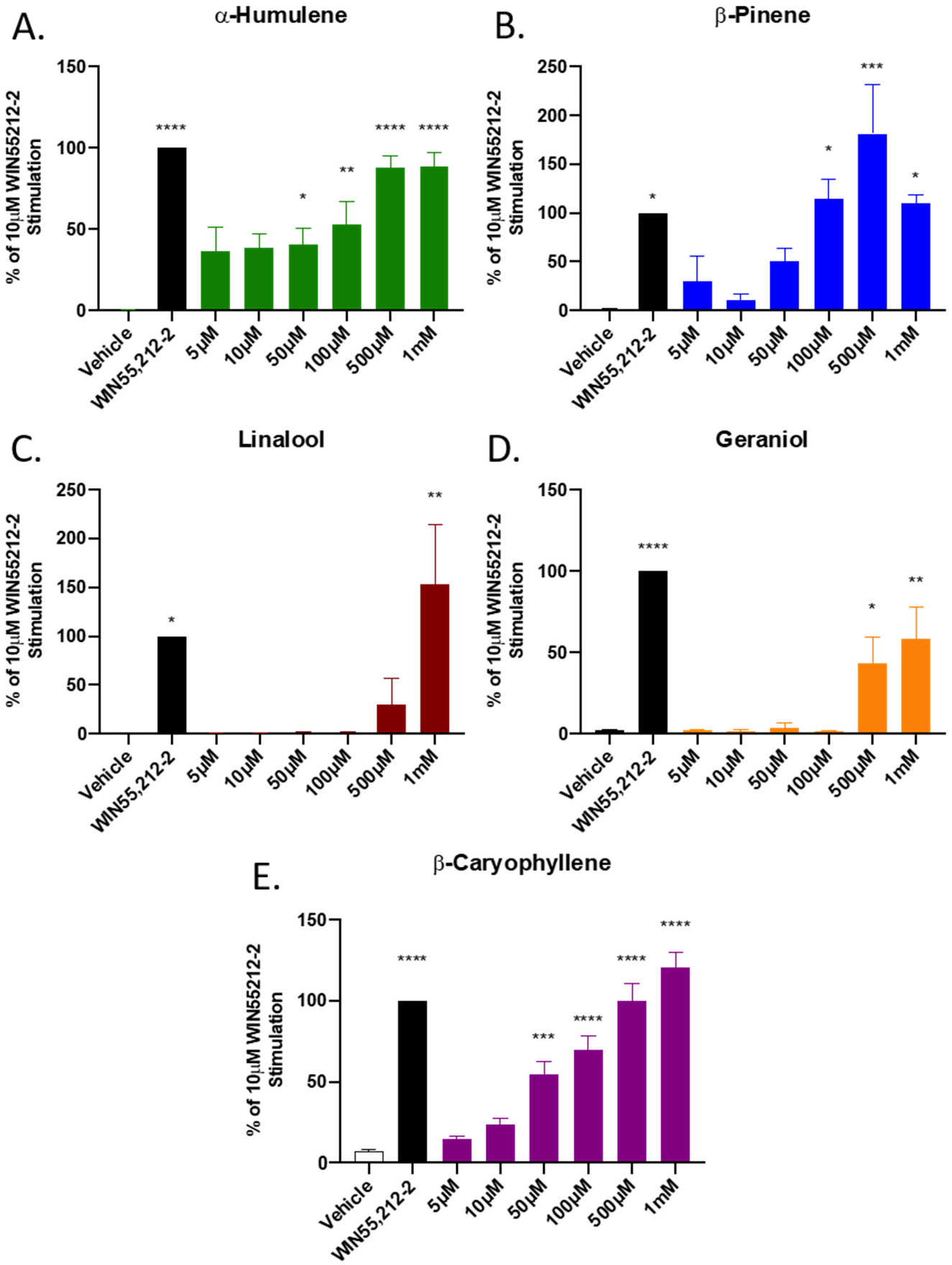
Terpene Treatment Activates the CB1 *In Vitro*. CB1-CHO cells were serum starved for 1 hr then treated with varying concentrations of **A)** α-Humulene, **B)** β-Pinene, **C)** Linalool, **D)** Geraniol, **E)** β-Caryophyllene, along with 10 μM WIN55,212-2 positive control or matched vehicle control, for 5 min. Lysates were then subjected to Western analysis and blotted for phospho-ERK and total-ERK (see Methods). Graph represents the quantified Western bands (pERK/tERK). Data expressed as a % of WIN55,212-2 stimulation (n=3 independent experiments). Statistics analyzed via one-way ANOVA, Dunnett’s *post hoc*; * p<0.05, ** p<0.01, *** p<0.001, **** p<0.0001, compared to vehicle stimulation.

**Figure 7:**
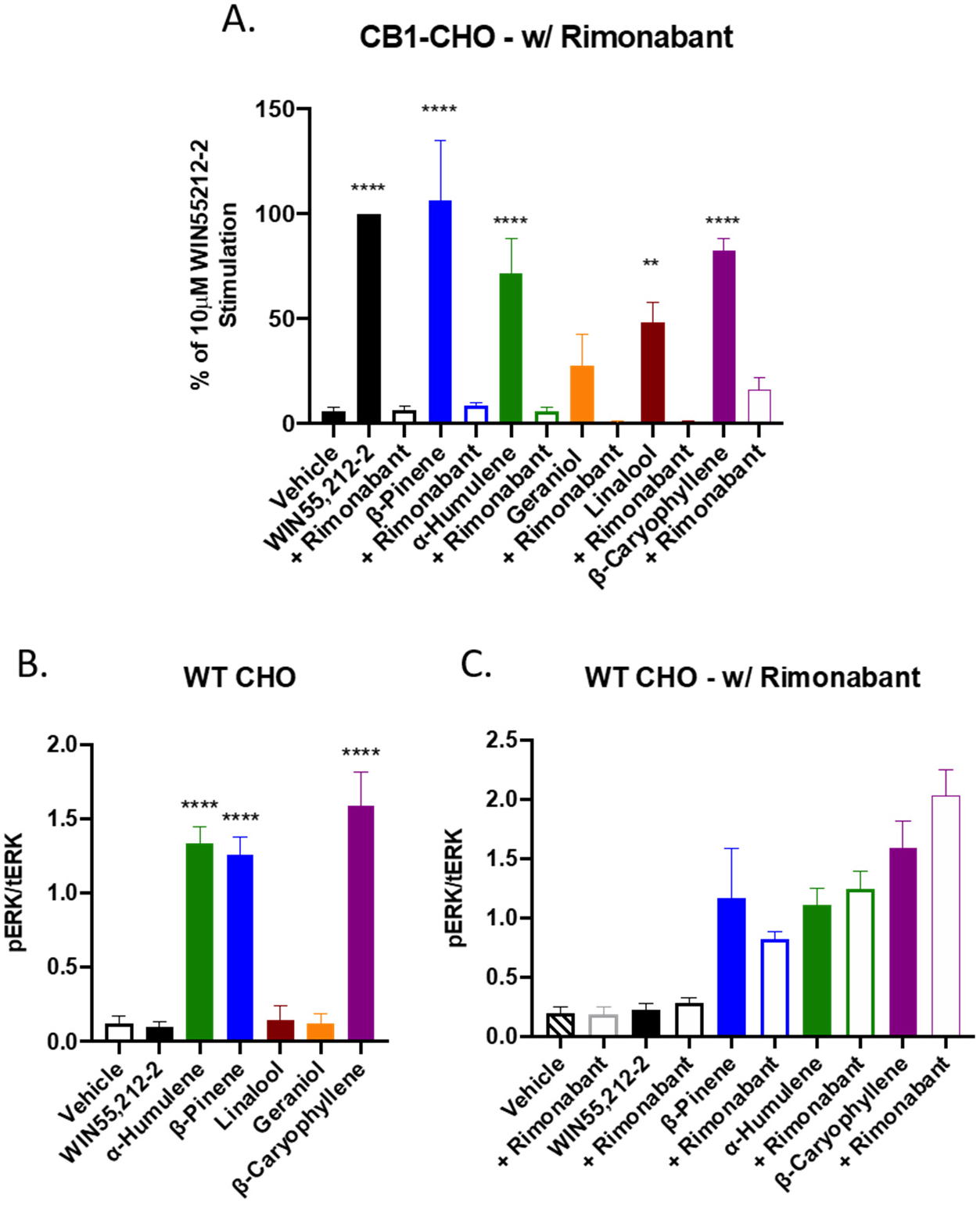
Terpenes Induce CB1-Dependent and Independent Signaling *In Vitro*. **A)** CB1-CHO cells were serum starved for 1 hr then pretreated with 10 μM rimonabant or vehicle for 5 min. Cells were then treated with 500 μM terpene, 10 μM WIN55,212-2, or matched vehicle, for 5 min. Lysates were then subjected to Western analysis and blotted for phospho-ERK and total-ERK (see Methods). Graph represents the quantitation of ERK phosphorylation induced by terpene and rimonabant combinations (pERK/tERK). Data expressed as a % of WIN55,212-2 stimulation (n=3 independent experiments for each). **B)** WT CHO cells were serum starved for 1 hr then treated with 500 μM terpene, 10 μM WIN55,212-2, or matched vehicle, for 5 min, and analyzed as above. **C)** WT CHO cells were serum starved for 1 hr, pretreated with 10 μM rimonabant or vehicle, then treated with 500 μM terpene, 10 μM WIN55,212-2, or matched vehicle, for 5 min, and analyzed as above. Data analyzed via one-way ANOVA, Dunnett’s *post hoc*; ** p<0.01, **** p<0.0001, compared to vehicle stimulation.

However, we found evidence that some terpenes act at other receptors or effectors to activate ERK *in vitro* when we screened the terpenes in WT-CHO cells presumably lacking any CB receptor **(Figures 7B, S10C)**. Linalool, Geraniol, and WIN55,212-2 did not cause ERK phosphorylation in WT CHO cells, however, α-Humulene, β-Pinene and β-Caryophyllene activated ERK in a rimonabant-insensitive manner **(Figures 7C, S10D-E)**. As rimonabant can act as an inverse agonist, potentially confounding the results above, we tested several concentrations of rimonabant against fetal bovine serum (FBS)-induced ERK activation **(Figure S11)**. Rimonabant did not significantly reduce ERK phosphorylation due to FBS treatment. This evidence suggests that each of the tested terpenes induced phosphorylation of ERK that is dependent on the CB1 receptor while some activated non-CB1 targets in these cells.

Of note, each of the terpenes tested also caused ERK phosphorylation in CB2-expressing cells **(Figure S12)**, suggesting they may interact with CB2 as already described for β-Caryophyllene [13]. Together, this evidence supports terpene poly-pharmacology, that can evoke behavioral and cellular changes via CB1-dependent and CB1-independent mechanisms (e.g. Adenosine A2a and CB2 from above).

We next followed up with a more comprehensive analysis of the pharmacological properties of each terpene at CB1. We first ran competition binding assays in CB1-CHO membrane preparations to determine whether each would compete for the orthosteric binding site against CP55,940 **(Figure 8A)**. As shown, WIN55,212-2 induced a typical concentration response-curve, fully competing CP55,940 away at higher concentrations. Of the terpenes assessed, Geraniol was the only one that displayed full competition. α-Humulene and β-Caryophyllene both displayed some competition and semi-biphasic properties. Linalool and β-Pinene displayed little to no competition. These results suggest both orthosteric and potential allosteric binding mechanisms for some of the different terpenes at CB1.

**Figure 8:**
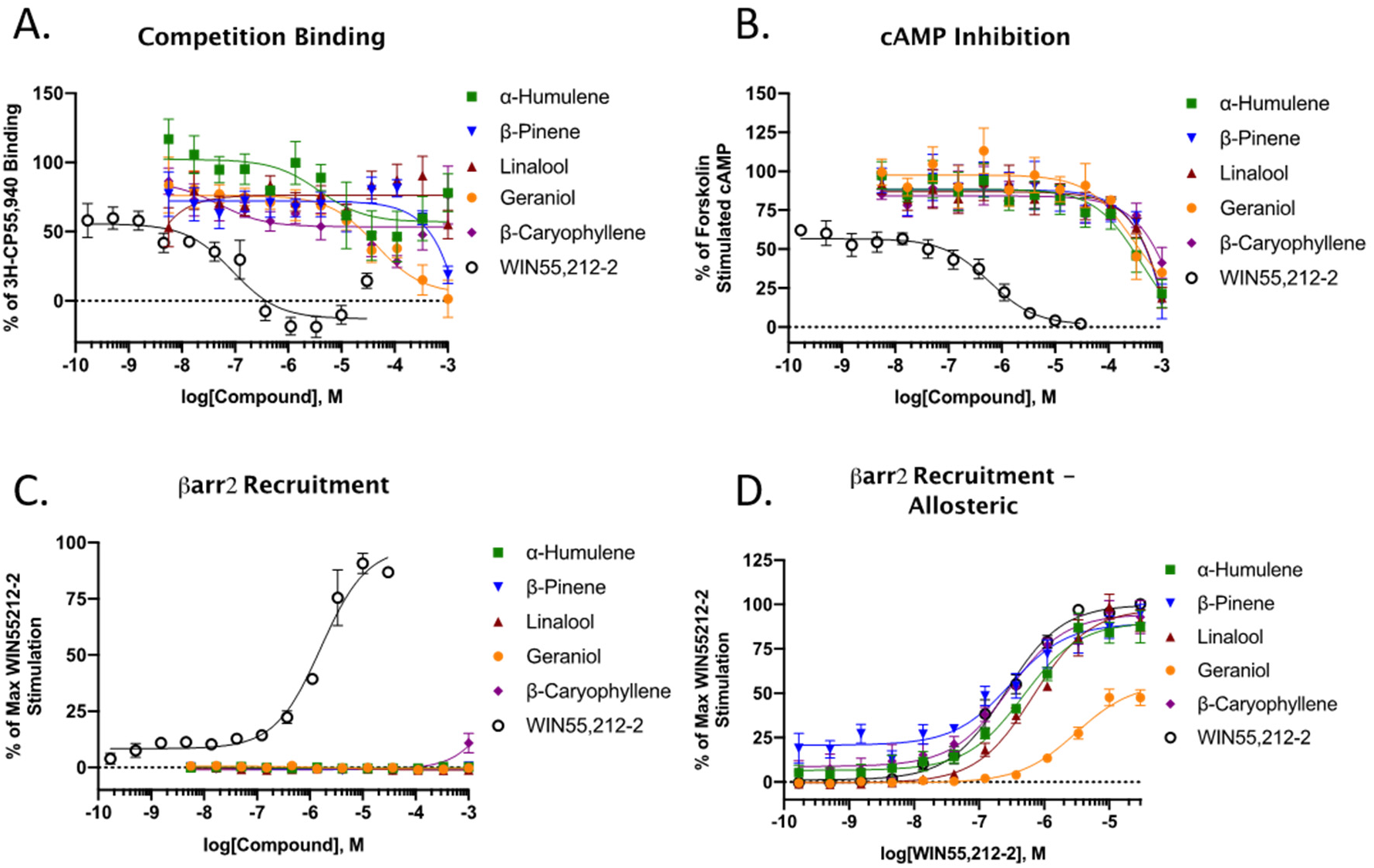
Binding and Functional Analysis of Terpenes at the CB1. **A)** CB1-CHO membranes were subjected to competition binding assay with terpenes and WIN55,212-2 against ^3^H-CP55,940. Data represents the mean ± SEM of specific binding (n=5 independent experiments). **B)** CB1-CHO cells were treated with varying concentrations of terpene or WIN55,212-2 for 30 min. The ability to inhibit forskolin-stimulated cAMP accumulation was measured accordingly (see Methods). Data represents the mean ± SEM of % of forskolin-stimulated cAMP (n=4 independent experiments). **C)** CB1-CHO-DX cells were treated with varying concentrations of compounds and βarrestin2 recruitment was assessed after 1.5 hr of treatment (see Methods). Data represents the mean ± SEM of max WIN55,212-2 recruitment (n=3 independent experiments). **D)** CB1-CHO-DX cells were pretreated with 500 μM terpene for 5 min, followed by varying concentrations of WIN55,212-2 for 1.5 hr (see Methods). Data represents the mean ± SEM of max WIN55,212-2 recruitment (n=3 independent experiments).

Following activation of the CB1 receptor, two downstream signaling pathways can be initiated; inhibition of adenylyl cyclase and β-arrestin2 recruitment. We first looked at the ability of the terpenes to inhibit adenylyl cyclase in CB1-CHO cells. Each of the terpenes tested demonstrated partial inhibition of forskolin-stimulated cAMP but only at high micromolar concentrations, while WIN55,212-2 displayed high potency and full efficacy **(Figure 8B)**. This effect by WIN55,212-2 and terpenes was partially reversed by rimonabant, except in the case of β-Pinene **(Figure S13A)**. Rimonabant had no effect on forskolin-stimulated cAMP formation by itself, suggesting no inverse agonism in this assay **(Figure S13B)**. These results suggest CB1-mediated downstream signaling induced by the terpenes.

When assessed in a βarrestin2 recruitment assay, terpenes displayed no agonism at any concentration tested, while the positive control WIN55,212-2 did **(Figure 8C)**. We next determined whether the terpenes may prevent or promote arrestin recruitment in an allosteric fashion and screened each terpene in combination with WIN55,212-2 **(Figure 8D)**. Only Geraniol displayed any presence of potential allosteric interaction and suggested it may be a negative allosteric modulator (NAM) or a biased partial agonist vs. WIN55,212, discussed below. When a full dose-range of Geraniol was tested against WIN55,212-2 it showed dose-dependent reductions in WIN55,212-2 recruitment **(Figure S13C)**.

## Discussion

As reviewed above, terpene compounds have been tested for their therapeutic properties in a number of studies, identifying anti-inflammatory, anti-nociceptive, anti-microbial, and similar beneficial properties [3, 4]. However very few studies have made any attempt to identify molecular targets and mechanisms for these compounds. Similarly, while terpenes and cannabinoids have been hypothesized to interact to produce an “entourage effect”, the few studies performed to date have shown no interaction [29, 30]. This study is thus the first to show that terpenes and cannabinoids can produce an additive effect when combined, providing support to the entourage effect hypothesis. This study is also the first to identify the CB1 and A2a receptors as terpene targets, and describe the role of these receptors in producing terpene cannabimimetic effects *in vivo*.

One question engendered by these findings is why our study found evidence for terpene/cannabinoid interactions while two other studies did not. There are several potential explanations. One study used an AtT20 cell line transfected with human CB1 or CB2 receptors, with a specific outcome of membrane potential change [29]. This study did not observe CB1/2 effects by terpenes while we did; however, the study was performed in a highly specific cell line with a single signaling output, membrane potential change. It is thus quite possible that the terpenes may not produce membrane potential changes via the CB1/2, just as we observed ERK and cAMP changes but not arrestin recruitment (**Figure 8**). The second study used a similar *in vitro* approach, finding no receptor binding or cAMP signaling of the terpenes tested [30]. However, this study used mixtures of terpenes, with no single terpene exceeding 50% of the mixture; they also used a maximum concentration of 10 μM, while we observed receptor activation at higher concentrations. These differences explain our observed results, and in addition, we used an *in vivo* model, which will capture a broader range of potential activity than an *in vitro* model with very specific outputs.

Another notable aspect of our study was the generally high concentrations of terpene needed to see activation. This was especially apparent in our *in vitro* studies, where >10 μM, or up to 500 μM depending on the terpene, was needed to see activation. Although high concentrations were required *in vitro*, this activation was still fully CB1-dependent, as it could be fully reversed by rimonabant treatment (**Figure 7A**). However, the doses needed *in vivo* were not as extreme, producing full responses in most assays for most terpenes at 200 mg/kg. This is consistent with the hypothesis that low potency multifunctional compounds may have benefits over selective single target compounds by inducing a significant systemic effect via multiple molecular effectors [31-34]. In this light, compounds, such as these terpenes, that have low potency at multiple receptor targets are likely to generate a substantial systemic effect (depending on the targets thereof). This idea is aligned with our *in vitro* data showing low potency effects and our *in vivo* data suggesting significant behavioral effects compared to a full CB1 agonist. Although it is clear that these selected terpenes can interact with the CB1 receptor to produce signaling, much more work is necessary to determine the mechanism of such action, and the collective group of terpene targets.

These observations have shown an important role for the CB1, among others, in mediating the effects of the test terpenes both *in vitro* and *in vivo*. Although appearing to be low potency agonists at CB1, there are alternative hypotheses for the mechanism of action based on these results. Two alternatives to direct CB1 agonism are 1) direct modulation of membrane dynamics, shifting CB1 activation equilibrium to favor the activated receptor; and 2) terpene modulation of endocannabinoid synthesis and/or degradation, which would then result in CB1 activation by these endocannabinoids. It has been suggested that membrane composition and dynamics heavily influence the cannabinoid receptor [35]. Indeed, this has also been described for other receptors [36] whereby membrane composition alters activation equilibrium. Terpenes are highly lipophilic in nature, and thus likely have direct interactions with the membrane environment, potentially also including membrane microdomains such as lipid rafts. If these interactions lead to a thermodynamically favored “active” CB1 receptor, the inverse agonism properties of rimonabant would still block signaling. In regards to the second point, it has been shown that CHO cells can participate in autocrine and paracrine signaling via endocannabinoid synthesis [37, 38], in these cases 2-arachadonylglycerol (2-AG). In our assays using CHO cells, it is thus a possibility that terpenes modulate the synthesis or degradation of 2-AG to stimulate CB1 signaling, which would be blocked by rimonabant. These alternative hypotheses are further supported by the generally weak competition binding observed at the CB1 by terpenes (**Figure 8A**). This is under current investigation in our laboratory.

Our studies suggest terpene poly-pharmacology, with multiple receptor targets, including CB1/2 and A2a. Other authors who tested some of these terpenes in the context of essential oils have detected glutamatergic [39], serotonergic [40], or dopaminergic [41] activities. Co-activation of such GPCR systems or ion channels with CB1 and/or CB2 receptors may explain why we observed differences between the terpenes for interaction with cannabinoid and the influence of CB1 and A2a on different tetrad behaviors. Thus, it seems that the observed effects produced by these terpenes are the complex outcomes of the activation or inhibition of multiple receptor systems.

This complex poly-pharmacology may provide a unique means to use terpenes to enhance cannabinoid or other therapies. In our findings, we see that all terpenes synergize with WIN55,212-2 to produce enhanced anti-nociception (**Figure 2**) while interacting variably with WIN55,212-2 in the other behaviors (**Figures 3-5**). In principle, this suggests that terpenes could be used to enhance the analgesic properties of cannabis/cannabinoid therapy, without worsening the side effects of cannabinoid treatment. Identifying specific terpene:cannabinoid combinations with a maximized therapeutic index for a particular disease state could provide a new means to improve human therapy with these drugs.

## Materials and Methods

### Materials

WIN55,212-2 (Tocris, #1038), α-Humulene (Sigma Aldrich, #53675), β-Pinene (Alfa Aesar, #A17818), Linalool (Alfa Aesar, #A14424), Geraniol (Alfa Aesar, #13736), and β-Caryophyllene (Cayman, #21572) were all prepared as stock solutions in DMSO. Working solutions were then diluted in 10% DMSO, 10% Tween-80, and 80% USP saline for injections. Rimonabant (Tocris, #0923) was made up in DMSO and then diluted to 20% DMSO, 10% Tween-80, and 70% USP saline for injections. Istradefyllene (Tocris, #5147) was made up in DMSO and then diluted to 20% DMSO, 10% Tween-80 and 70% saline for injections. Morphine sulfate pentahydrate (from the NIDA Drug Supply Program) was dissolved in USP saline. Vehicle injections were matched accordingly as a control in each experiment. All solutions were made immediately before use without long-term storage. For *in vitro* experiments, 100 mM stocks of terpenes, and 10 mM stocks of all other compounds, were made up in DMSO. DMSO concentrations in assays were maintained at 1% or lower DMSO. Vehicle controls were matched accordingly.

### Animals

All experiments were performed on male and female CD-1 (a.k.a. ICR) mice obtained from Charles River in age-matched cohorts of 5-6 weeks of age. All mice were recovered for at least 5 days after shipping prior to experimentation, and housed no more than 5 per cage. The animals were maintained in an AAALAC-accredited vivarium at the University of Arizona in temperature and humidity-controlled rooms on a 12-hour light/dark cycle. Standard chow and water were available *ad libitum*. For all behavioral experiments, mice were brought to the testing room in their home cages for at least 30 minutes prior to the experiment for acclimation. Mice were randomized to treatment group, and the experimenter was blinded to treatment group by the delivery of coded drug vials. The groups were not decoded until all data had been acquired. All experiments were approved by the University of Arizona IACUC, and were carried out in accord with the standards of the NIH Guide for the Care and Use of Laboratory Animals. We also adhered to the guidelines of ARRIVE and the *British Journal of Pharmacology*; no adverse events were noted for any of the animals.

### Tail Flick

Antinociception was tested using the tail flick thermal latency test. Mice were baselined by gently restraining the animal and lowering the distal portion of the tail into a water bath set to 52°C or 47°C where stated, and the latency to flick the tail recorded with a stopwatch, with a 10 second cutoff. Mice were then injected with compounds intraperitoneally (i.p.). Thermal latency was then assessed over a time course of two hours, or at a single time-point as noted. For assays using rimonabant, rimonabant was injected 10 min prior to terpene or morphine injection. For assays using terpene/WIN55,212-2 co-treatment, WIN55,212-2 was co-administered with terpene in the same solution.

### Catalepsy

Potential induction of catalepsy was assessed using the ring test as described [42]. Mice were baselined, injected with compound, and then assessed with the ring test at 15 min post-injection. Data was represented as the percentage (%) of time in a “cataleptic” immobile state over the 5 minute observation period.

### Open Field Test

The open field test was used to determine potential hypolocomotive properties induced by terpene and cannabinoid treatment. The open field box was a white box with an open top and black floor. The tracking area was 30cm x 28cm. Mice were baselined by placing the mouse in the center of the box and recording from a video camera ∼1.5 m above for a period of 5 min. Mice were then injected with compound and placed back in the open field box 10min-15min post-injection. In follow-up experiments, mice were not baselined, injected with compound, and placed in the open field box from 10min-15min. The total distance traveled and mobile time of mice were analyzed using AnyMaze software.

### Hypothermia

Changes in core temperature were assessed using a lubricated rectal thermometer placed 1.0 cm into the mouse rectum. Temperature was assessed before treatment and 30 min post-treatment.

### Cell Culture

WT-CHO cells were obtained from ATCC (#CCL-61). CHO cells stably expressing the human CB1 receptor were obtained from PerkinElmer (#ES-110-C) and utilized for western blots (CB1-CHO-C2) or binding assays (CB1-CHO-C3). HitHunter (CB1-CHO-cAMP) and PathHunter (CB-CHO-βarr2) assays were performed on respective CB1-CHO lines, and were a kind gift from Dr. Robert Laprairie [43]. CHO cells stably expressing cloned CB2 (CB2-CHO) were produced by electroporation with the human CB2 N-3xHA tag cDNA (GeneCopoeia). A stable expressing population was selected for with 500 μg/mL G418. All cells were grown on 10 cm dishes in DMEM/F-12 50/50 mix w/ L-glutamine and 15 mM HEPES (Corning) containing 10% heat-inactivated fetal bovine serum, 100 units/mL penicillin, 100 units/mL streptomycin, and 400 μg/mL G418 (CB1-CHO-C2, CB1-CHO-C3, CB2-CHO); 800 μg/mL G418 (CB1-CHO-cAMP); 800 μg/mL G418 and 300 μg/mL hygromycin B (CB1-CHO-βarr2). WT CHO were grown as stated above without selection antibiotic. Cells were incubated in a humidified incubator at 37°C with 5% CO2 and were sub-cultured every 2-3 days.

### Cell Treatments for ERK

Cells were plated in 6-well plates 24 hr prior to the start of the experiments. Cells were serum starved for 1 hr then treated for 5 min with compound. In antagonist experiments, cells were pretreated for 5 min before agonist treatment for 5 min. Media was then aspirated, cells were placed on ice, washed with ice-cold PBS once, and lysis buffer (20mM Tris-HCl, pH 7.4, 150 mM NaCl, 2 mM EDTA, 0.1% SDS, 1% Nonidet P-40, 0.25% deoxycholate, 1mM sodium orthovanadate, 1mM PMSF, 1mM sodium fluoride, and a protease inhibitor cocktail (EMD Millipore)), was added. Cells were scraped and collected. Lysates were vortexed then centrifuged at 13k rpm at 4°C for 10min. Lysates were then processed for Western blots or stored at -80°C.

### Electrophoresis and Western Blotting

Cell lysate protein was quantified with a modified BCA protein assay using manufacturers protocol (Bio-Rad). Samples (20-30 μg) were ran on precast gels (10% Bis-Tris, Bolt brand, from ThermoFisher). Gels were wet-transferred to nitrocellulose membranes at 30 V for at least 60 min at 4°C. Blots were blocked with 5% non-fat dry milk in TBS for 30 min, washed 3X for 5 min with TBS + 0.1% Tween-20 (TBST), and incubated with primary antibody overnight at 4°C. Primary antibody: pERK and tERK (Cell Signaling) at 1:1000 dilution in 5% BSA in TBST. Blots were then washed 3x with TBST then incubated with secondary antibodies for 90 min at room temperature. Secondary antibodies: Goat anti-Rabbit 800W and Goat anti-Mouse 680 diluted at 1:10,000 and 1:20,000 (Licor), respectively, in 5% non-fat milk in TBST. Blots were washed 3X with TBST and 1X with TBS, then imaged on a Licor Odyssey Fc or Azure Sapphire. Bands were analyzed using Image-J and reported as pERK/tERK normalized to the standard, or simply as pERK/tERK.

### Radioligand Binding

Radioligand binding was performed as previously reported [44]. Briefly, CB1-CHO-C3 cells were homogenized in 50mM Tris-HCl containing 1mM PMSF, then centrifuged at 30,000g for 30 min at 4°C. The resultant pellet was resuspended in the same buffer. 30-40 μg of membrane protein was incubated with varying concentrations of compound and 0.3 nM [^3^H]-CP55,940 (PerkinElmer) for 90 min at room temperature. Data reported as the % of specific [^3^H]-CP55,940 binding.

### PathHunter Assay

CB1-CHO-βarr2 cells were plated in 384-well plates (5,000 cells/well) in Opti-MEM with 1% FBS. The following day cells were treated with varying concentrations of ligand for 1.5 hr. In allosteric assays, cells were pre-treated with the first compound for 5 min. Manufacturers protocol was then followed. Luminescence was read on a CLARIOstar plate reader. Data reported as the % of maximum recruitment by WIN55,212-2 reference standard.

### HitHunter Assay

CB1-CHO-cAMP cells were plated in 384-well plates (5,000 cells/well) in Opti-MEM with 1% FBS. The following day, the media was removed and replaced with PBS. Cells were then treated with varying concentrations of compounds in solution containing 20 μM forskolin (Enzo) for 30 min. In antagonist experiments cells were pre-treated for 5 min. Following agonist treatment, the manufacturers protocol was followed. Data reported as the % of forskolin stimulated cAMP.

### Data Analysis and Statistics

All data was analyzed using GraphPad Prism 8. For behavioral studies graphs show combined male and female data. When qualitative sex differences were observed, males and females were separated for analysis. For dose-response curves, non-linear fit curves were generated using Prism. The behavioral data was all reported in raw values, without normalization. Statistical analysis details are noted in the Figure Legends. For RM 2 Way ANOVA, the Geisser-Greenhouse correction was used to account for a potential lack of sphericity of the data, permitting valid RM ANOVA. ANOVA *post hoc* tests were only performed when ANOVA F values indicated a significant difference, and there was homogeneity of variance (permitting parametric analysis). In all cases, significance was defined as *p* < 0.05. Statistical analysis for *in vivo* experiments was only performed where N ≥ 5. Our experimental design and analysis further comply with the recommendations and requirements of the *British Journal of Pharmacology* [45].

## Abbreviations

2-AG: 2-Arachidonoylglycerol
A2a: Adenosine A2a Receptor
CBD: Cannabidiol
CB1/2: Cannabinoid Receptor Type 1/2
ERK: Extracellular Signal-Regulated Kinase
FBS: Fetal Bovine Serum
NAM: Negative Allosteric Modulator
THC: Δ9-Tetrahydrocannabinol
TBST: Tris-Buffered Saline with Tween

## Acknowledgments

The authors would like to acknowledge Drs. Tally Largent-Milnes and Todd Vanderah from the University of Arizona for sharing their behavioral equipment and expertise. This study was funded by institutional funds from the University of Arizona. JMS has an equity stake in *Botanical Results, LLC*, a local cannabidiol company; no company products or interests were tested in this study. The authors have no other relevant conflicts of interest to declare.

## Data Availability

The data that support the findings of this study are available from the corresponding author upon reasonable request.

