## Supplementary Figures for "*Cannabis sativa* Terpenes are Cannabimimetic and Provide Support for the Entourage Effect Hypothesis"

**This file includes:**
Figures S1 to S13

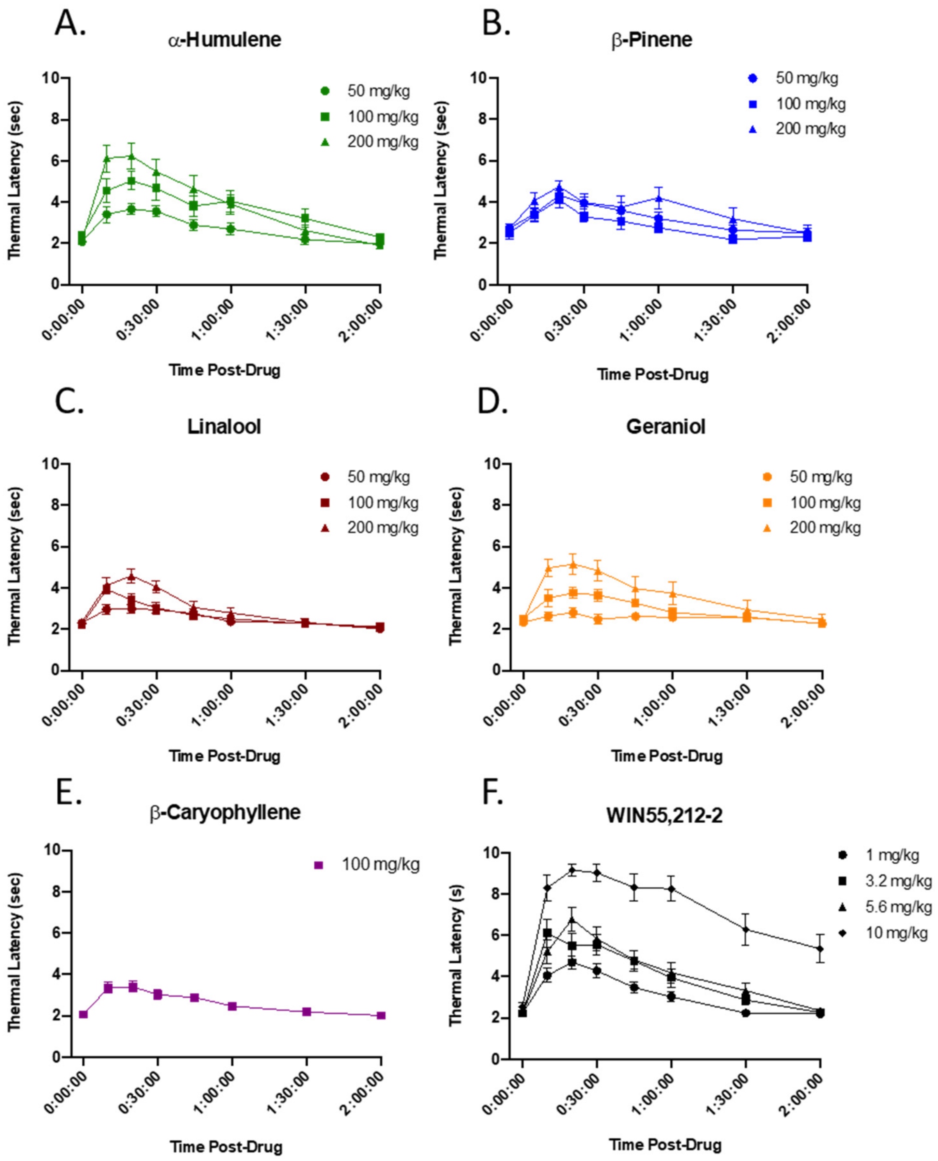


**Figure S1: Terpenes Are Antinociceptive in the Tail Flick Assay.** Mice were treated with varying doses of terpene (**A-E**) or WIN55,212-2 (**F**), *intraperitoneal* (*i.p.*), and assessed in the tail flick thermal latency test over a period of 2 hours. Data represents the mean ± SEM of tail flick latency in seconds (n=10/group).

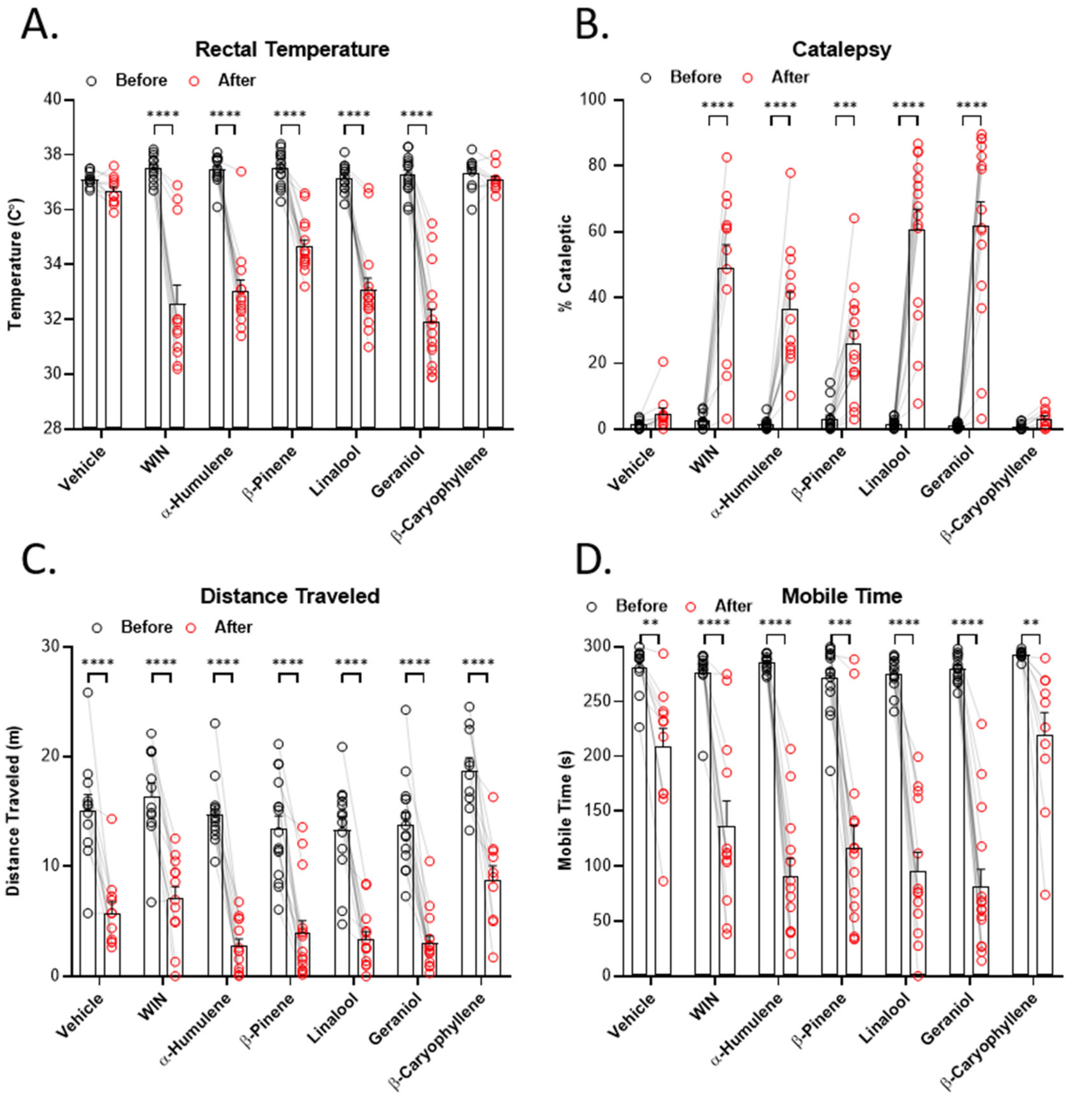


**Figure S2: Terpenes Induce Hypothermia, Catalepsy, and Hypolocomotion. A)** Mice were tested for temperature at baseline and 30 min after *i.p.* injection with 200 mg/kg terpene, 5.6 mg/kg WIN55,212-2, or matched vehicle. Data represents the mean ± SEM of temperature (n=10-15/group). **B)** Each mouse was baselined in the ring test for 5 min, then again at 15 min after *i.p.* injection with 200 mg/kg terpene, 5.6 mg/kg WIN55,212-2, or matched vehicle. Data represents the mean ± SEM of % catalepsy (n=10-15/group). **C) and D)** Mice were injected with 200 mg/kg terpene, 5.6 mg/kg WIN55,212-2, or matched vehicle and then tested in the open field test after 10 min, for 5min, and analyzed using ANYmaze software. Data represents the mean ± SEM of mobile time in seconds (**C**) or distance traveled in meters (**D**) (N=10-13/group). Statistics analyzed via RM two-way ANOVA, Dunnett's *post hoc*; bracket = $p<0.05$ vs. each baseline measurement.

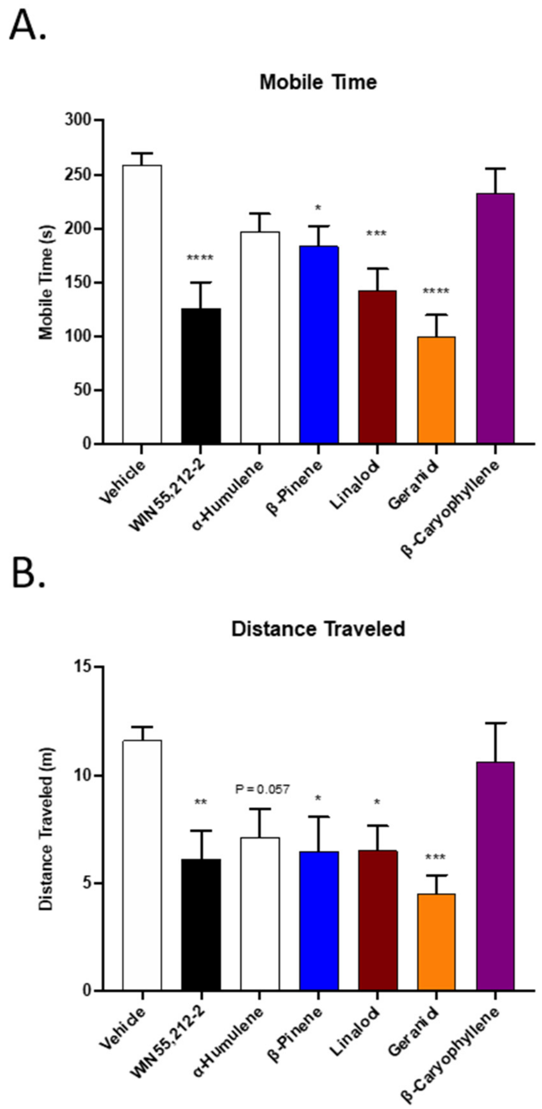


**Figure S3: Terpenes Induce Measures of Hypolocomotion.** Mice were baselined in the open field test for 5 min then injected with 200 mg/kg terpene, 5.6 mg/kg WIN55,212-2, or matched vehicle, *i.p.*. After 10 min mice were then placed back into the open field box for a 5 min test. Measures of **A)** distance traveled and **B)** mobile time were analyzed using ANYmaze software. Data represents the mean ± SEM of distance traveled (**A**) and mobile time (**B**) (n=10-15/group). Statistics analyzed via one-way ANOVA, Sidak's *post hoc*; ** $p<0.01$, *** $p<0.001$, **** $p<0.0001$ vs. Vehicle group.

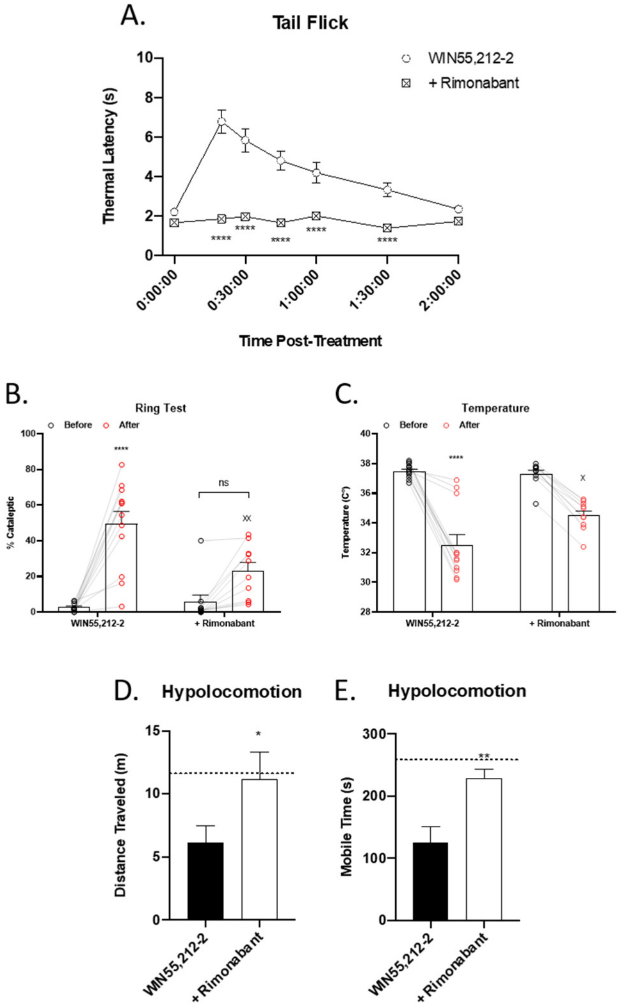


**Figure S4: WIN55,212-2 Induced Tetrad Effects Are Mediated by the CB1 Receptor.** Mice were treated with 5.6 mg/kg WIN55,212-2 or after pretreatment with 10 mg/kg rimonabant, *i.p.*. **A)** Mice were then assessed in the tail flick test over 2 hr. Data represents the mean ± SEM of tail flick latency (n=10/group). Statistics analyzed via two-way ANOVA, Dunnet's *post hoc*; **** $p<0.0001$ compared to WIN55,212-2 alone. **B)** Mice were baselined in the ring test for 5min, injected as above, and after 15 min, mice were tested in the ring test again for 5 min. Data represents the mean ± SEM of % catalepsy (n=10-12/group). Statistics analyzed via two-way ANOVA, Tukey's *post hoc*; **** $p<0.0001$ compared to baseline, xx $p<0.01$, compared to WIN55,212-2 post-treatment. **C)** Mice were baselined for temperature, injected as above, and after 30 min, temperature was assessed again. Data represents the mean ± SEM of temperature (n=10-12/group). Statistics analyzed via two-way ANOVA, Tukey's *post hoc*; **** $p<0.0001$ compared to baseline, x $p<0.05$ compared to WIN55,212-2 post-treatment. **D) and E)** Mice were baselined in the open field test for 5 min, injected as above, and after 10 min mice were then placed back into the open field box for a 5 min test. Data represents the mean ± SEM of distance traveled (**D**) and mobile time (**E**) (n=10-13/group). Statistics analyzed via unpaired 2-tailed *t* test; * $p<0.05$, ** $p<0.01$ compared to WIN55,212-2 alone. Dotted line denotes vehicle levels for reference.

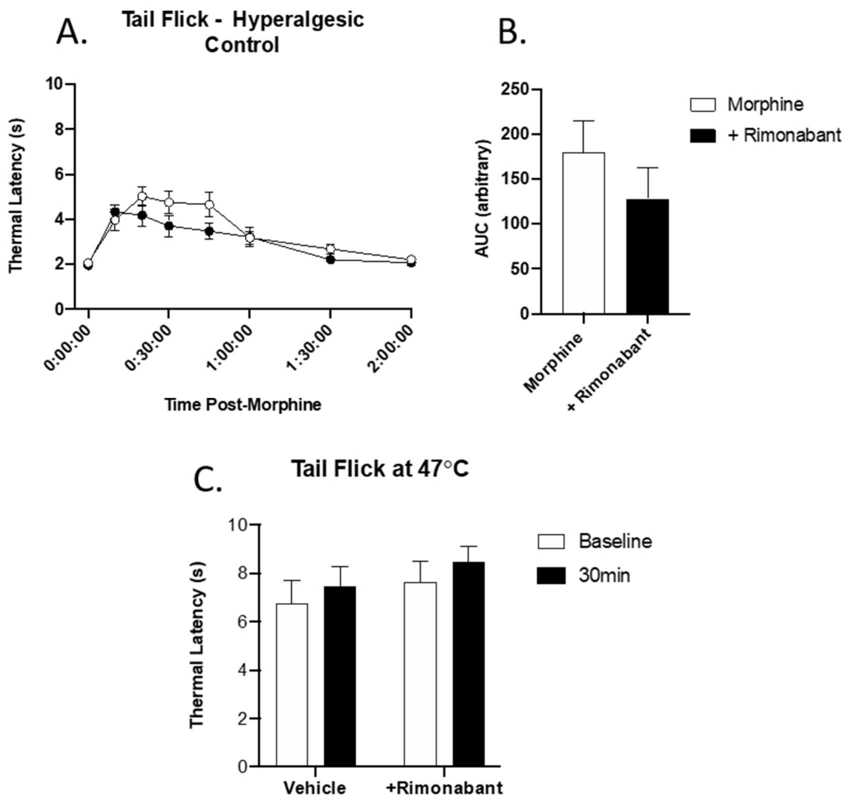


**Figure S5: Rimonabant Does Not Act as An Inverse Agonist in the Tail Flick Assay. A)** Mice were treated with 5.6 mg/kg morphine or after pretreatment with 10 mg/kg rimonabant, *i.p.*. Mice were then assessed in the tail flick test over 2 hr. Data represents the mean ± SEM of tail flick latency (n=10/group). No statistical differences observed via two-way ANOVA. **B)** Area under the curve analysis of **A**. No statistical differences observed via t-test. **C)** Mice were baselined at 47ºC, then injected with 10 mg/kg rimonabant or matched vehicle. After 30 min mice were baselined again. Data represents the mean ± SEM of tail flick latency (n=10/group). No statistical differences observed via two-way ANOVA.

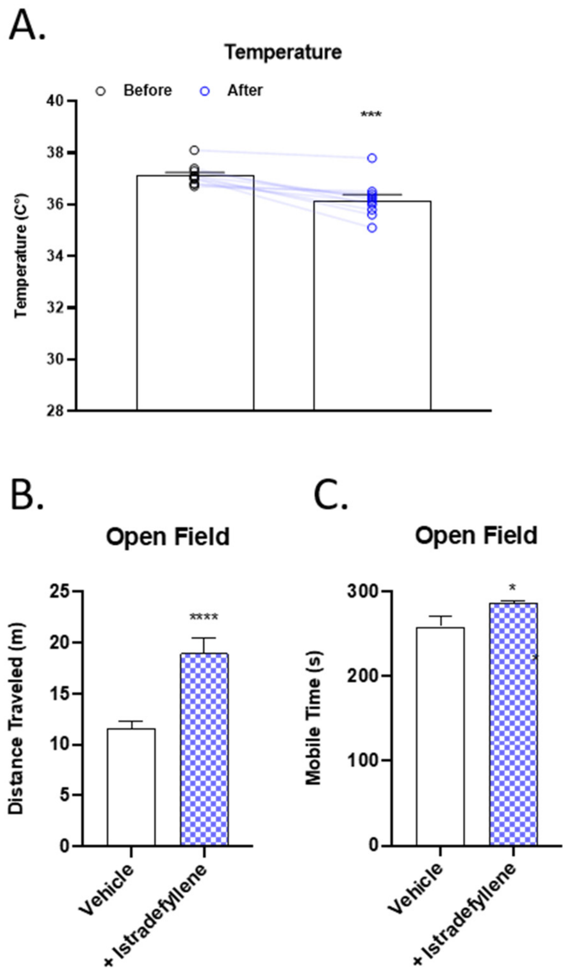


**Figure S6: Istradefyllene Treatment Causes Hypothermia and Hyperlocomotion. A)** Mice were baselined for temperature, then injected with 10 mg/kg istradefyllene, *i.p*. After 30 min, temperature was assessed again. Data represents the mean ± SEM of temperature (n=10). Statistics analyzed via two tailed paired t-test. *** p<0.001 compared to baseline. **B) and C)** Mice were injected with 10 mg/kg istradefyllene or vehicle, *i.p.*. After 10min mice were then placed back into the open field box for a 5 min test. Measures of **B)** distance traveled and **C)** mobile time were analyzed using ANYmaze software. Data represents the mean ± SEM of distance traveled (**B**) and mobile time (**C**) (n=10-12/group). Statistics analyzed via unpaired two tailed t-test. * p<0.05, **** p<0.0001, compared to vehicle.

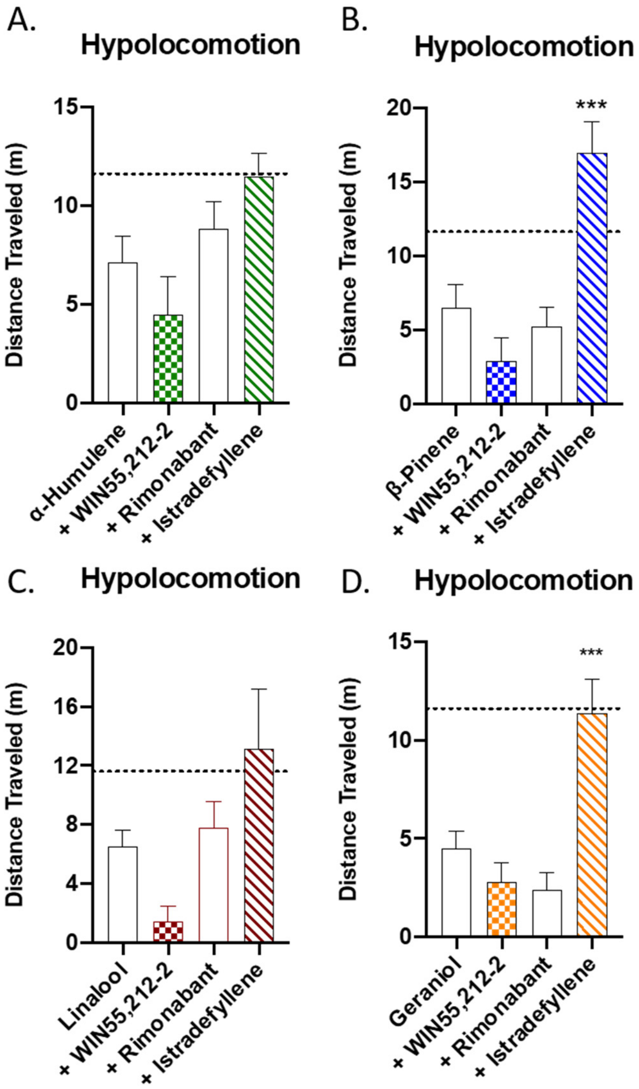


**Figure S7: Terpene Induced Hypolocomotion is Partially Mediated by A2a and is Additive with Cannabinoid.** Mice were injected with 200 mg/kg terpene alone, combined with 5.6 mg/kg WIN55,212-2, or after pretreatment with 10 mg/kg rimonabant or 10 mg/kg istradefyllene, *i.p.*. After 10 min mice were then placed back into the open field box for a 5 min test. Measures of distance traveled were analyzed using ANYmaze software. **A)** α-Humulene, **B)** β-Pinene, **C)** Linalool and **D)** Geraniol. Data represents the mean ± SEM of distance traveled (n=10-20/group). Statistics analyzed via one-way ANOVA, Dunnett's *post hoc*; *** $p<0.001$ compared to terpene alone. Dotted line denotes vehicle levels of distance traveled for reference.

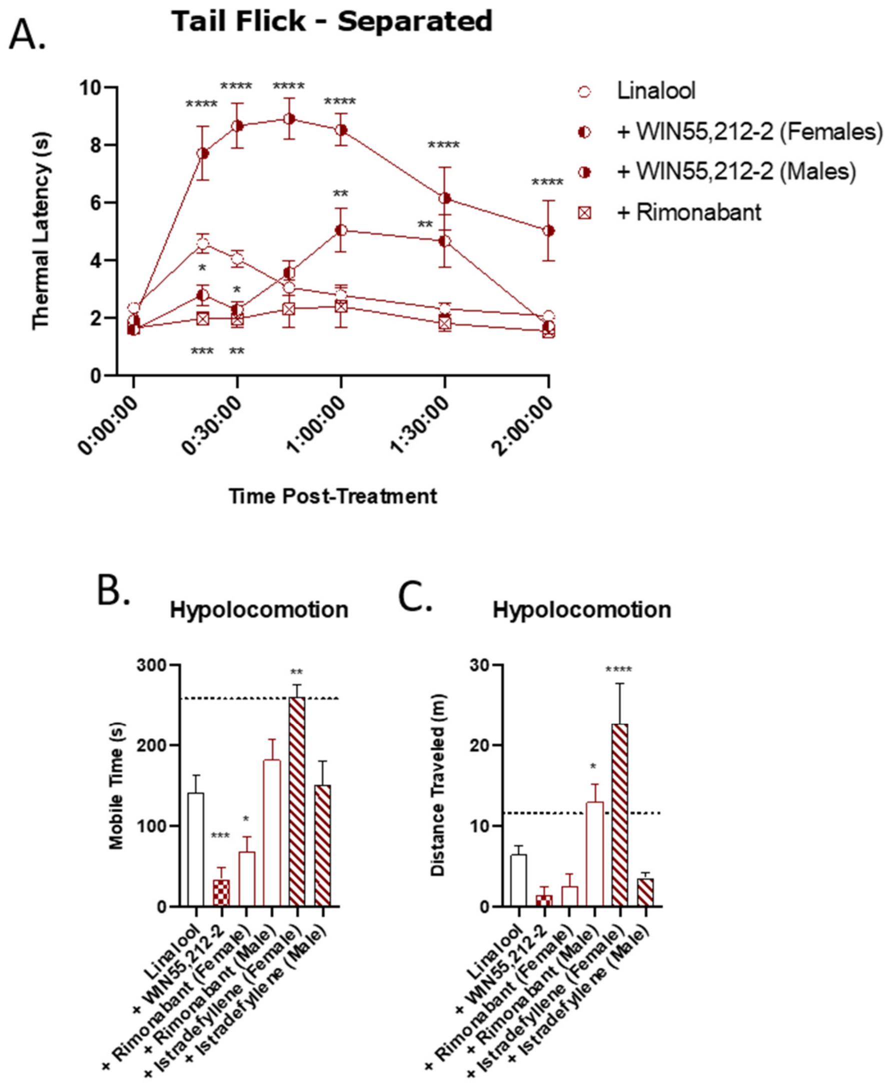


**Figure S8: Sex-Differences in Linalool Mechanism of Action.** Mice were tested as described in **Figure 1, 3, 4 and S4** and separated where sex differences were qualitatively observed. **A)** Linalool modulation of tail flick is differentially modulated by WIN55,212-2 treatment. Data represents the mean ± SEM of tail flick latency (n=7-15/group). Statistics analyzed via two-way ANOVA, Dunnett's *post hoc*; *p<0.05, ** p<0.01, *** p<0.001, **** p<0.0001, compared to Linalool alone. **B)** Females display istradefyllene-sensitive cataleptic responses. Data represents the mean ± SEM of % catalepsy (n=4-10/group). Statistics analyzed via two-way ANOVA, Tukey's *post hoc*; **** p<0.0001 compared to baseline, xxx p<0.001, xxxx p<0.0001, compared to Linalool post-treatment. **C) and D)** Hypolocomotive behavior, as described above, separated by sex. Data represents the mean ± SEM of distance traveled (**C**) and mobile time (**D**) (n=5-17/group). Statistics analyzed via one-way ANOVA, Dunnett's *post hoc*; * p<0.05, ** p<0.01, *** p<0.001, **** p<0.0001, compared to Linalool alone.

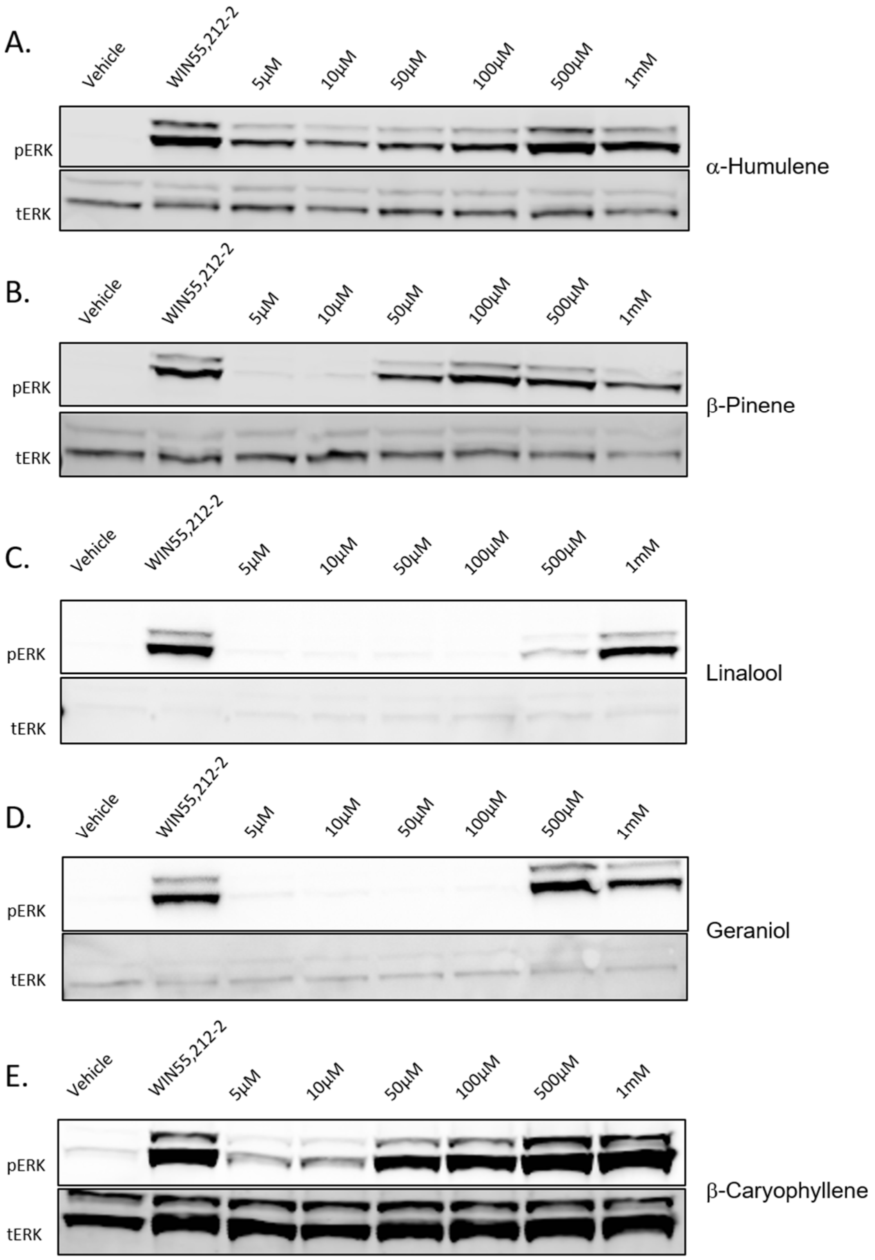


**Figure S9: Terpene Treatment Activates the CB1 *In Vitro*.** CB1-CHO cells were serum starved for 1 hr then treated with varying concentrations of **A)** α-Humulene, **B)** β-Pinene, **C)** Linalool, **D)** Geraniol and **E)** β-Caryophyllene, along with 10 µM WIN55,212-2 or matched vehicle controls, for 5 min. Representative blots shown for data found in **Figure 6**.

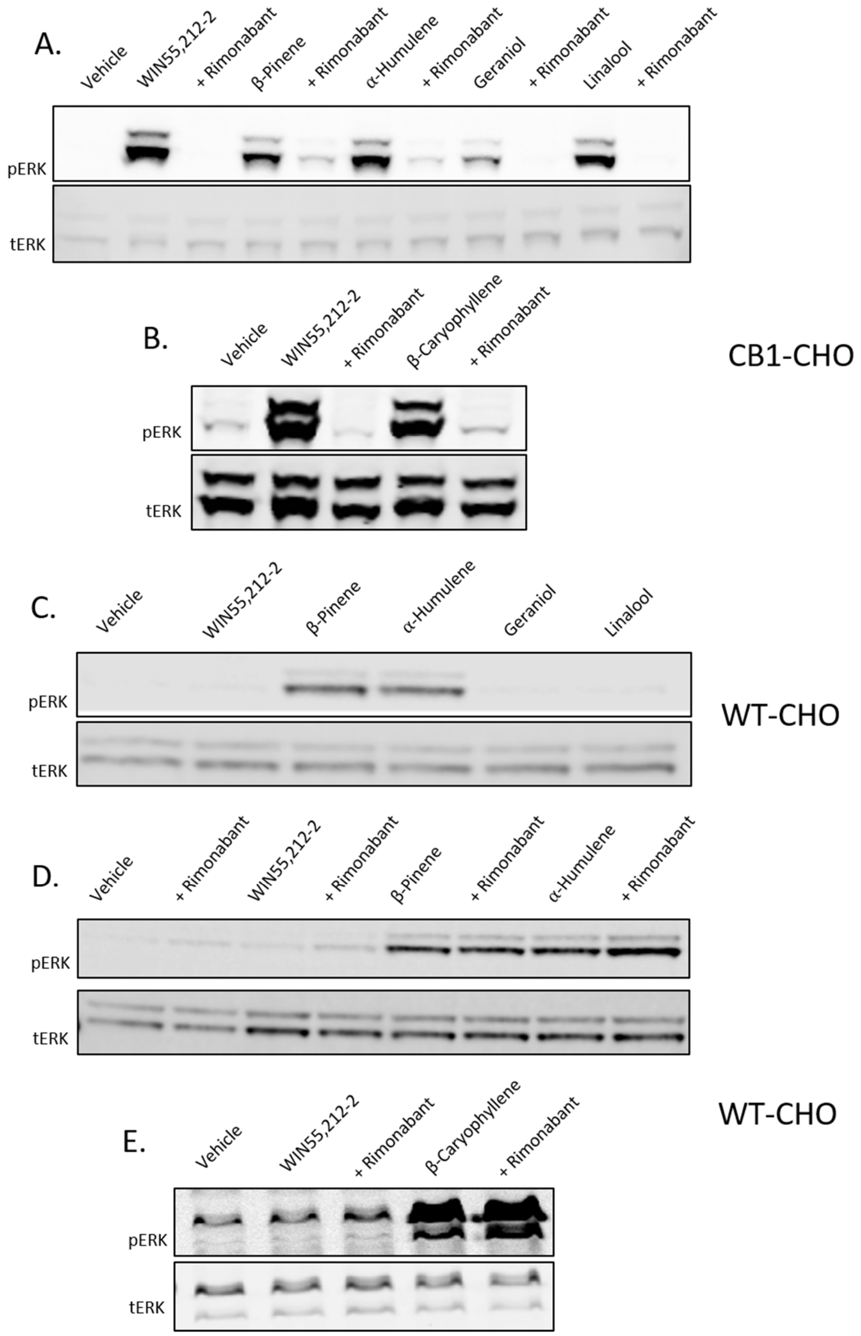


**Figure S10. Terpenes Induce CB1-Dependent and Independent Signaling *In Vitro*.** Representative blots shown for data in **Figure 7. A and B)** CB1-CHO cells were serum starved for 1 hr then pretreated with 10 μM rimonabant or vehicle for 5 min. Cells were then treated with 500 μM terpene, 10 μM WIN55,212-2, or matched vehicle, for 5 min. **C)** WT CHO cells were serum starved for 1 hr then treated with 500 μM terpene, 10 μM WIN55,212-2, or matched vehicle, for 5 min. **D) and E)** WT CHO cells were serum starved for 1 hr, pretreated with 10 μM rimonabant or vehicle, then treated with 500 μM terpene, 10 μM WIN55,212-2, or matched vehicle, for 5 min.

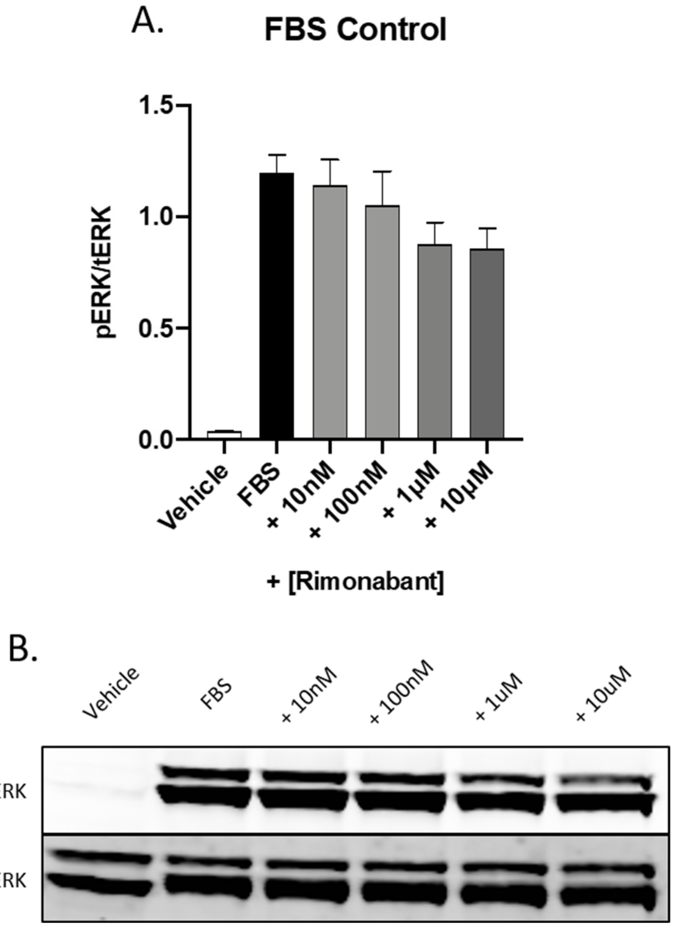


**Figure S11: Rimonabant Does Not Block FBS-Stimulated ERK Phosphorylation in CB1-CHO Cells.** CB1-CHO cells were serum starved for 1 hr, pretreated with varying concentrations of rimonabant or vehicle, and then treated with 10% FBS for 5 min. Lysates were then subjected to Western analysis and blotted for phospho-ERK and total-ERK (see Methods). **A)** Western quantitation of ERK phosphorylation. Data expressed as phospho-ERK/total-ERK (n=3 independent experiments). Statistics analyzed via one-way ANOVA showed no differences when compared to FBS only stimulation. **B)** Representative Western blot image from the data in **A**.

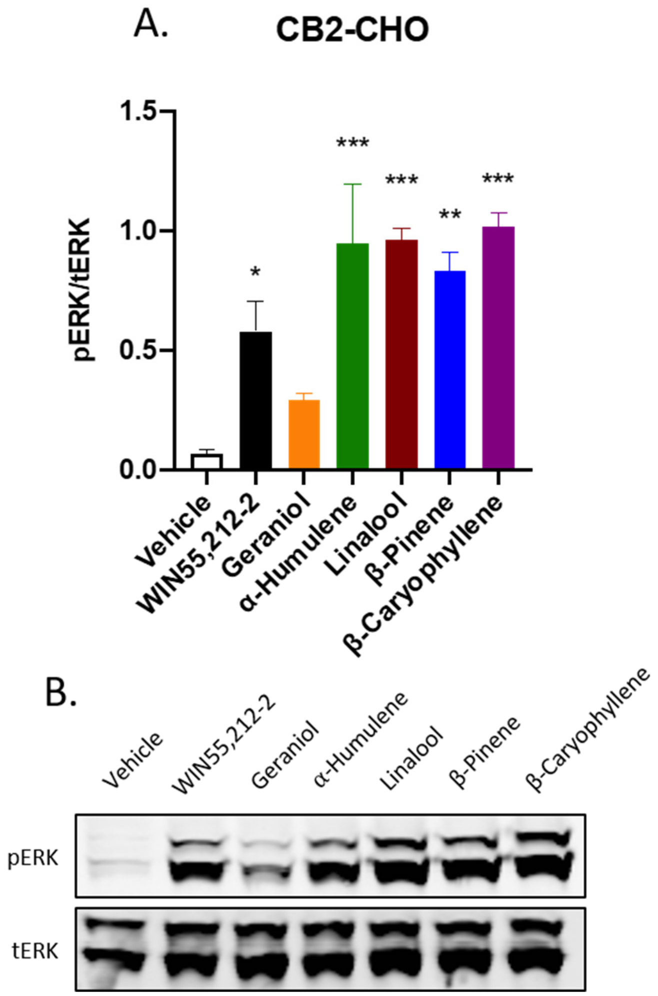


**Figure S12: Terpenes Induce ERK Phosphorylation in CB2-CHO Cells.** CB2-CHO cells were serum starved for 1 hr then treated with 500 µM terpene, 10 µM WIN55,212-2, or vehicle, for 5 min. Lysates were then subjected to Western analysis and blotted for phospho-ERK and total-ERK (see Methods). **A)** Western quantitation of ERK phosphorylation induced by terpenes in CB2-CHO cells. Data expressed as phospho-ERK/total-ERK (n=3 independent experiments). Statistics analyzed via one-way ANOVA, Dunnett's *post hoc*; * $p<0.05$, ** $p<0.01$, *** $p<0.001$, compared to vehicle stimulation. **B)** Representative Western blot image shown for the data in **A**.

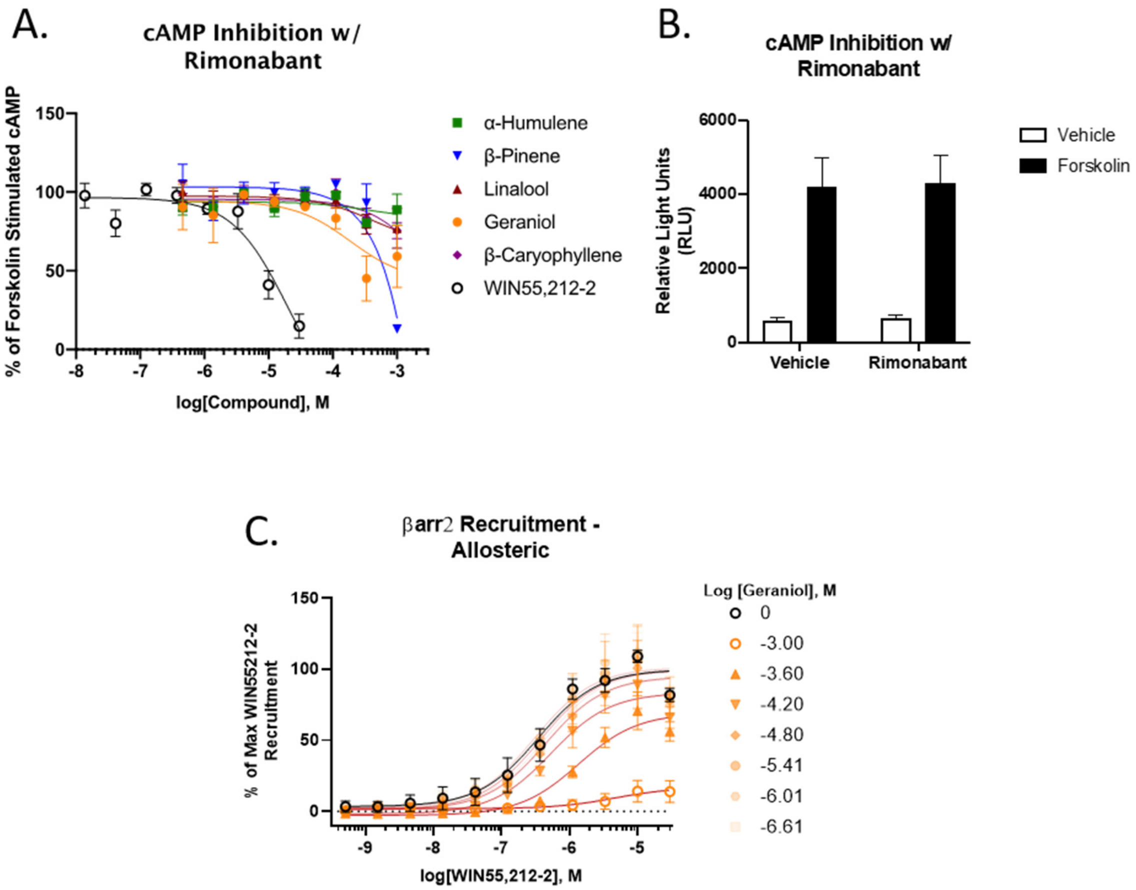


**Figure S13: Binding and Functional Analysis of Terpenes at the CB1. A)** CB1-CHO cells were pretreated with 10 μM rimonabant, then treated with varying concentrations of terpene or WIN55,212-2 for 30 min. The ability to inhibit forskolin-stimulated cAMP accumulation was then measured (see Methods). Data represents the mean ± SEM of % of forskolin-stimulated cAMP (n=4 independent experiments). **B)** Vehicle and forskolin data from **A**, depicting lack of inverse agonism by rimonabant. Data represents the mean ± SEM of the RLU (n=4 independent experiments). **C)** CB1-CHO-DX cells were pretreated with varying concentrations of Geraniol for 5 min, followed by varying concentrations of WIN55,212-2 for 1.5 hr (see Methods). Data represents the mean ± SEM of max WIN55,212-2 recruitment (n=3 independent experiments).
